# Localized modulation of DNA supercoiling, triggered by the *Shigella* anti-silencer VirB, is sufficient to relieve H-NS-mediated silencing

**DOI:** 10.1101/2023.01.09.523335

**Authors:** Michael A. Picker, Monika M. A. Karney, Taylor M. Gerson, Alexander D. Karabachev, Juan C. Duhart, Joy A. McKenna, Helen J. Wing

**Affiliations:** School of Life Sciences, University of Nevada Las Vegas, Las Vegas, NV 89154-4004, USA

## Abstract

In Bacteria, nucleoid structuring proteins govern nucleoid dynamics and regulate transcription. In *Shigella spp*., at ≤ 30 °C, the histone-like nucleoid structuring protein (H-NS) transcriptionally silences many genes on the large virulence plasmid. Upon a switch to 37 °C, VirB, a DNA binding protein and key transcriptional regulator of *Shigella* virulence, is produced. VirB functions to counter H-NS-mediated silencing in a process called transcriptional anti-silencing. Here, we show that VirB mediates a loss of negative DNA supercoils from our plasmid-borne, VirB-regulated *PicsP-lacZ* reporter, *in vivo*. The changes are not caused by a VirB-dependent increase in transcription, nor do they require the presence of H-NS. Instead, the VirB-dependent change in DNA supercoiling requires the interaction of VirB with its DNA binding site, a critical first step in VirB-dependent gene regulation. Using two complementary approaches, we show that VirB:DNA interactions *in vitro* introduce positive supercoils in plasmid DNA. Subsequently, by exploiting transcription-coupled DNA supercoiling, we reveal that a localized loss of negative supercoils is sufficient to alleviate H-NS-mediated transcriptional silencing, independently of VirB. Together, our findings provide novel insight into VirB, a central regulator of *Shigella* virulence and more broadly, a molecular mechanism that offsets H-NS-dependent silencing of transcription in bacteria.

## INTRODUCTION

Dynamic genome management is a critical process in all living cells because it underpins DNA replication, influences transcription and controls cellular physiology (1,2). Bacterial genomes, once considered naked, are, in fact, shaped and managed by nucleoid structuring proteins (NSPs). These proteins act as “genome sentinels”, controlling the sequestration or liberation of parts of the genome, so that genes lie silent or become active when accessed by the cell’s molecular machinery (3–6). The molecular events that trigger the remodelling of NSPs remain poorly understood and yet these processes are functionally equivalent to epigenetic processes that regulate gene expression in eukaryotic cells. As we face increasing antibiotic resistance in a wide swath of bacterial pathogens (7), improved knowledge of these critical regulatory processes may reveal novel bacterial targets that can be exploited by new therapeutics. As such, transcriptional silencing and anti-silencing has become a burgeoning field in the realm of bacterial gene regulation (8–13).

In Bacteria, the histone-like nucleoid structuring protein (H-NS) is widely distributed among the Proteobacteria (renamed Pseudomonadota) but functional homologs exist in other phylogenetic groupings, including the Firmicutes (renamed Bacillota) (14–18). H-NS preferentially binds AT-rich DNA (19–21), which is a common feature of horizontally acquired loci (22). In this capacity, H-NS silences spurious promoters and horizontally acquired genes, aiding the retention of newly acquired DNA (12,22–26), which often includes key virulence genes in important bacterial pathogens (15,27–31). Although a high affinity binding site for H-NS has been proposed (19,32), AT-rich DNA tracts produce narrow minor groove widths that primarily govern the DNA binding preference of H-NS (33). H-NS binding to DNA is followed by its oligomerization along the helix into regions with lower binding affinities (32,34), leading to the formation of large H-NS:DNA complexes. Two H-NS:DNA complexes have been described, depending on external conditions: H-NS nucleoprotein filaments that coat long, contiguous stretches of DNA, and H-NS bridging complexes that bring two discrete regions together through direct H-NS-H-NS interactions (35–39). Both of these nucleoprotein complexes have been implicated in the silencing of virulence genes in *Shigella* species, where the complexes either occlude promoter elements or trap RNA polymerase (RNAP) in loops that hinder transcription (reviewed by (13)).

Transcriptional anti-silencing describes the unveiling or remodeling of H-NS coated DNA to allow it to become accessible for transcription (13). Anti-silencing may be triggered by a change in environmental parameters, like osmolarity, pH, and temperature, which alter DNA topology (40,41). Alternatively, DNA binding proteins known as anti-silencers are required to trigger this process. Anti-silencers are a diverse group of DNA binding proteins (examples include Ler, LeuO, RovA, SlyA, VirB and AraC family members), which has led to the proposal that a variety of ‘ad hoc’ solutions offset the effects of H-NS (13). However, it is possible that this cohort of unrelated proteins exhibit shared mechanisms of transcriptional anti-silencing. Thus, detailed mechanistic studies of anti-silencing proteins and their shared regulatory processes may significantly broaden our general understanding of these key proteins and the vital role they serve in bacteria.

In *Shigella* species., the etiological agents of bacillary dysentery (42), the transcriptional anti-silencing protein VirB is encoded by the large virulence plasmid (43). Once made (44), VirB binds to its recognition site and alleviates H-NS mediated silencing of virulence plasmid loci (45–48), controlling around 50 virulence genes (49). This regulatory effect is essential for *Shigella* virulence, as *virB* mutants are avirulent (50–52). The interplay between VirB and H-NS has been investigated at four virulence plasmid loci, *icsP* (45–47), *icsB* (48), ospZ (53) and *ospD1* (54), but, arguably, mostly extensively at the *icsP* locus. Although the classical view of bacterial transcription is that promoter proximal events modulate levels of transcription initiation (55), this view is overly simplistic when considering transcriptional silencing and anti-silencing (56). Indeed, at the *icsP* locus, the region required for H-NS mediated silencing is located between 900 and 400 base pair upstream of the major transcription start site (TSS;+1) as depicted in Figure 1A (46). Furthermore, the required VirB binding site is located over 1 kb upstream (centred at −1137; Figure 1A) (46, 47,57). Strikingly, if a LacI protein is engineered to dock between the H-NS and VirB regulatory regions, VirB is unable to counteract H-NS silencing (46) suggesting that the intervening sequence is key in determining the VirB regulatory effect. Although experimental evidence indicates that VirB spreads along the DNA at the *icsP* locus toward the region bound by H-NS (46), the relevance of this and mechanistic details of how this ultimately remodels the H-NS silencing complex remains unclear.

**Figure 1.**
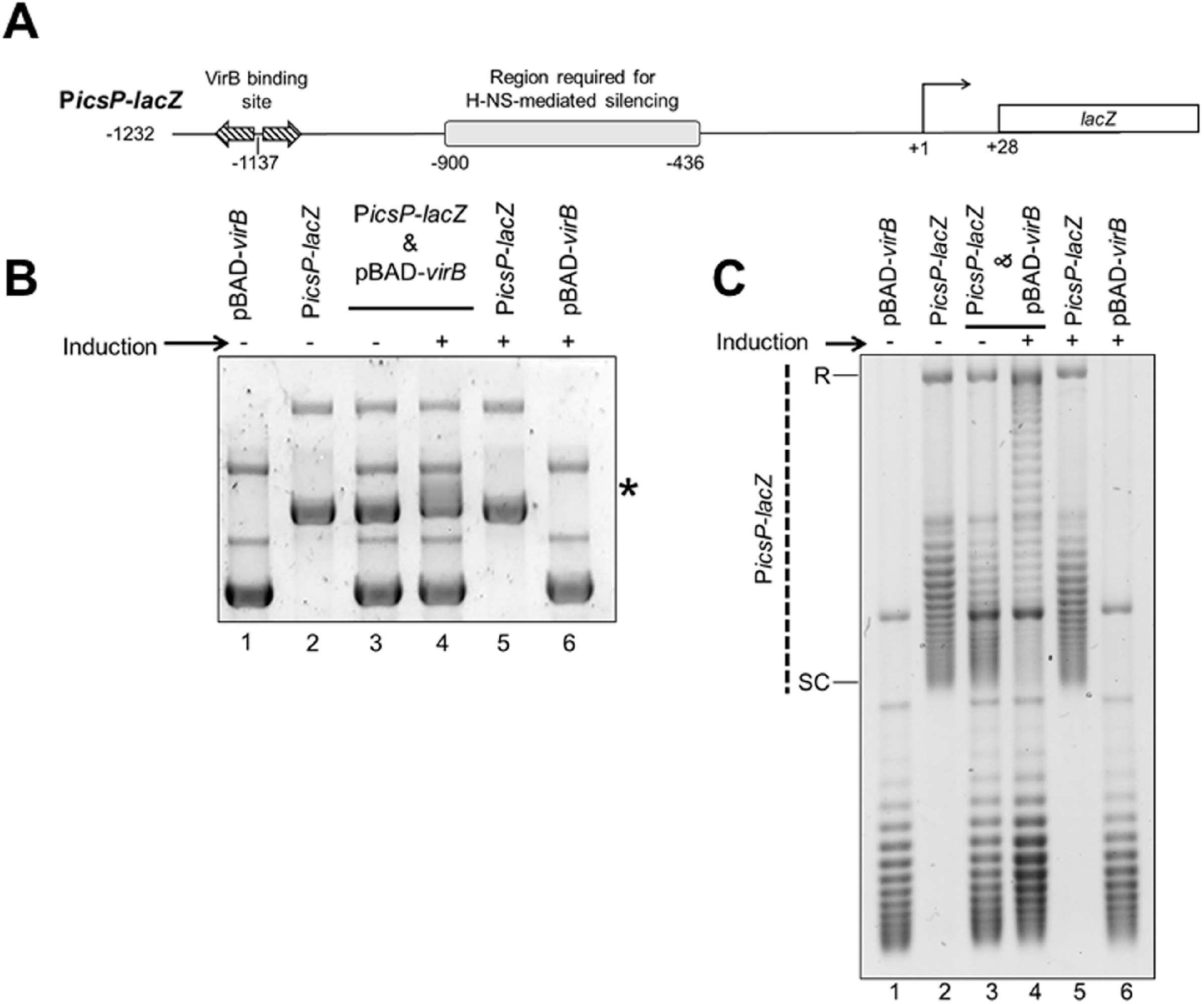
VirB triggers a change in electrophoretic mobility and topoisomer distribution of the *PicsP-lacZ* reporter plasmid. A) Schematic of VirB-dependent *PicsP-lacZ* carried by the reporter plasmid, pHJW20, used throughout this study (not drawn to scale). Previously identified regulatory elements are indicated (39, 40,48). Note, only primary TSS is shown (89). B) Electrophoretic mobility of wild-type *PicsP-lacZ* reporter isolated from *E. coli* DH10B in the presence or absence of *virB* induction during growth. *, denotes position of the change in electrophoretic mobility of *PicsP-lacZ*. This gel was run in the absence of ethidium bromide and stained afterwards. C) Analysis of *PicsP-lacZ* topoisomer distribution of samples from (A) with an agarose gel containing chloroquine (2.5 μg ml^-1^) [R=relaxed and SC=supercoiled]. Quantification of key lanes (3 & 4) on gels shown in B and C is provided in Figure S6 A-D.

Curiously, VirB does not belong to a family of classical transcription factors but rather is a distantly related member of the ParB superfamily of DNA partitioning proteins (48,58–60). Although VirB is not involved in plasmid segregation (60–62), we reasoned that its evolutionary lineage may provide important insight into its role as an anti-silencer. ParB proteins bind to their DNA recognition site, and spread along DNA forming long protein filaments (63,64), a process that causes changes in DNA supercoiling (65–67). More recently, some members of the ParB family have been shown to spread along DNA by forming a sliding clamp in a CTP-dependent manner (68–70). While VirB is distantly related to these proteins, VirB recognizes a ParB-like DNA binding site (46–48,57,58,60) and forms high order oligomers, both *in vitro* and *in vivo* (58,59). Furthermore the DNA binding and spreading activities of VirB are critical for transcriptional anti-silencing of virulence plasmid genes (46,57–59,71). In light of these findings, we hypothesized that VirB:DNA interactions *in vivo* & *in vitro* trigger changes in DNA topology that alleviate transcriptional silencing of the *icsP* promoter mediated by H-NS. Thus, the overarching goal of this study was to gain mechanistic insight into the bacterial process of transcriptional anti-silencing by VirB, which would likely inform our understanding of mechanisms used by other bacterial anti-silencing proteins.

## MATERIAL AND METHODS

### Bacterial strains, plasmids, and media

The bacterial strains and plasmids used in this study are listed in Table S1. *Escherichia coli* and *Shigella flexneri* cultures were routinely grown at 37°C on either LB agar [*E. coli;* LB broth with 1.5% agar (wt. vol^-1^)], or Congo Red plates [*S. flexneri;* Tryptic soy broth containing 0.01% (wt. vol^-1^) Congo Red], respectively. Congo Red positive phenotypes were used to verify that the large virulence plasmid of *Shigella* had been maintained, since it is prone to loss. Single, isolated colonies were grown overnight in LB broth with aeration at either 30°C (*S. flexneri; to* stabilize the virulence plasmid; {Schuch, 1997 #55}) or 37°C (*E. coli*). Overnight cultures were diluted 1:100 in fresh LB broth and allowed to grow for 5 h at 37°C. Where appropriate, antibiotics were added at the following final concentrations: ampicillin, 100 μg ml^-1^ and chloramphenicol, 25 μg ml^-1^.

### Strain and plasmid construction

The DH10B *hns* mutant derivative was constructed using a modified one-step method (72). Briefly, the *hns*::Kn^r^ locus was PCR amplified from MC4100 *hns* mutant (73) using primers W428 and W429 and the amplicon was electroporated into wild-type DH10B strain expressing the λ RED recombinase system (pKD46) [grown at 30°C in SOC medium (72)]. Recipients of the *hns*::Kn^r^ locus were selected for on LB agar containing kanamycin (25 μg ml^-1^), and the new strain was verified by sequencing using primers W428 and W429. To test the regulatory phenotype of the new DH10B *hns*::Kn^r^ strain, *icsP* promoter activity was measured in DH10B and the isogenic *hns* mutant using a β-galactosidase assay as previously described (45). Activity of the *icsP* promoter was significantly higher in the DH10B *hns*::Kn^r^ background compared to wild-type DH10B (data not shown), consistent with previous studies (45) using the same *hns* mutant locus in *E. coli* MC4100.

The plasmids used in this study are listed in Table S1. For all experiments, the VirB-dependent *PicsP-lacZ* reporter pHJW20, (pACYC184 derivative; (45)), or its derivatives were used. All plasmid constructs were verified by Sanger dideoxy sequencing. The sequence of all primers and DNA duplexes used in this work are listed in Table S2.

For the construction of pDivergent (pMK63) and pParallel (pMK64), a multi-step cloning procedure was used. Briefly, an insert (derived from G-block #13; IDT DNA) carrying 102 bp of the φ10 promoter sequence (identical to the sequence found in pTF1 (ATCC 77397)) transcriptionally fused to *gfp-2* (74) and followed by five *rrnB* T1 terminators (5xT) (75), was created (flanked by *PstI* and *Pvu*II (5’ end) and *Pvu*II and *Hin*dIII (3’ end); sequence provided in Table S1). This DNA insert was digested with *Pst*I and *Hin*dIII and ligate to pBlueScript II KS+ digested with the same restriction enzymes. To create constructs with *Pφ10-gfp-2-5xT* in two orientations, the resulting plasmid was digested with *Pvu*II and the restriction fragment containing the insert was ligated to pBlueScript II KS+ digested with *Eco*RV. This generated pMK34 and pMK35. These two plasmids were digested with *Pst*I and *XhoI* (located on the pBlueScript II KS vector), and each insert was ligated to pHJW20 digested with *Pst*I and *SalI*, to place the *Pφ10-gfp-2-5xT* insert in one of two orientations upstream of the *PicsP-lacZ* reporter. This generated the two plasmids used in this work pDivergent (pMK63) and pParallel (pMK64).

### *In vivo* topoisomer analysis

#### Plasmid Extraction

For *in vivo* supercoiling assays in *E. coli*, the *PicsP-lacZ* reporter or its derivatives, and the pBAD-*virB* expression plasmid or its derivatives, were introduced either alone or in combination into DH10B or its isogenic derivative *hns*::Kn^r^ mutant or into MG1655 and its isogenic derivative DPB923 (*topA* mutant) (Coli Genetic Stock Center). DH10B containing *PicsP-lacZ* (or derivatives) and inducible pBAD-*virB*, or DH10B with the single plasmid controls, were grown overnight, and with 0.2% (wt. vol^-1^) D-glucose (76) if the strains contained pBAD-*virB*. The following day, duplicate cultures were seeded from overnight cultures by diluting 1:100 in LB broth and these were grown for 3 h at 37°C. The expression of *virB* was induced in half of the cultures with the addition of 0.2% (wt. vol^-1^) L-arabinose, and all cultures were grown for 2 h more at 37°C. Cells were then harvested, normalised to cell density (OD_600_ nm), and processed using the PureYield Plasmid Miniprep System according to the manufacturer’s directions (Promega, Cat. No. A1222).

For *in vivo* supercoiling assays in *Shigella*, the *PicsP-lacZ* reporter was introduced into the wild-type *S. flexneri* strain 2457T or the isogenic *virB* mutant AWY3 strain. Overnight cultures of *Shigella* were subcultured for 5 h at 37°C with aeration. Cultures were normalised to cell density and DNA was isolated using the Plasmid Maxi Kit (Qiagen, Cat. No. 12163) according to manufacturer’s directions except that all steps and reagents, except Buffer P2, were held at 4°C and the DNA was eluted in TE buffer (10 mM Tris-HCl, pH 7.5 and 1 mM, EDTA, pH 8.0). These changes helped prevent excessive nicking and smearing of the DNA, as reported previously (77).

#### 1D Chloroquine Gel Electrophoresis

This protocol was adapted from Dorman:Choloroquine gel analysis, 2008 (OpenWetWare) and LeBlanc and Clark, 2009 (78,79). To analyse the topoisomer distribution of isolated DNA samples, routinely 500 ng of DNA was electrophoresed through a 2x TBE [178 mM Tris base, 178 mM Borate, 4 mM EDTA (pH 8.0)], 0.7% (wt. vol^-1^) agarose gel, for 16 - 24 h in the dark at 2.0 - 2.5 V cm^-1^ in the presence of chloroquine, unless otherwise noted. Routinely, a final concentration of 2.5 μg ml^-1^ was used in both gel and buffer, but an elevated concentration (10 μg ml^-1^) was used determine whether supercoiling changes caused by VirB were positive or negative (Figure S2). Note that DNA isolated from the DH10B *hns* mutant in the presence or absence of *virB* induction (Figure 3B) was instead electrophoresed through a 1% agarose gel at 4.7 V cm^-1^ for 1.5 h with chloroquine. After electrophoresis, gels containing chloroquine were destained in sterile distilled water, changing the water every 30 - 60 min. After destaining, gels were then stained with ethidium bromide, imaged using a UVP Biospectrum 410 Imaging System and resulting images were manipulated using Visionworks™ LS Image Acquisition.

**Figure 2.**
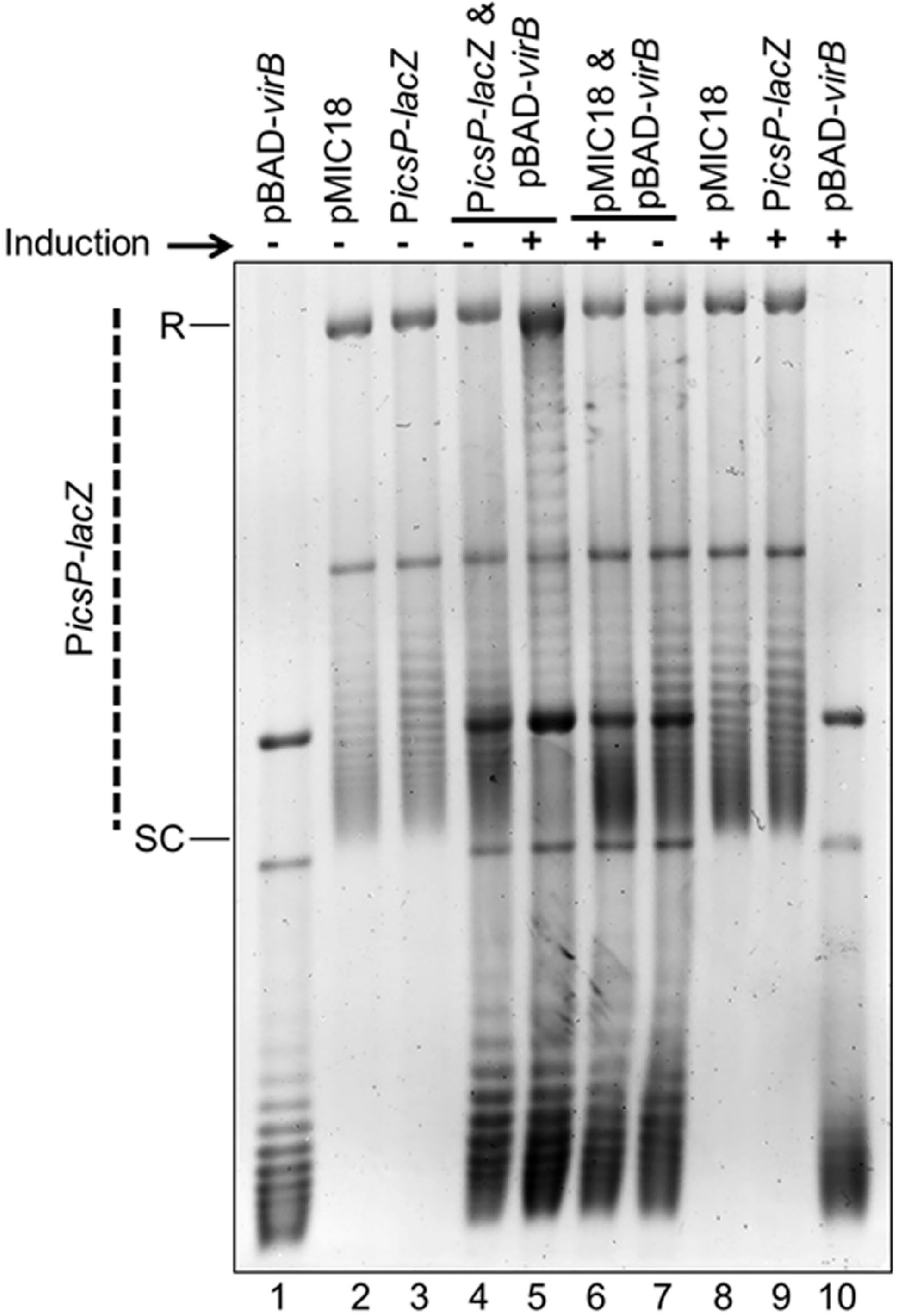
The VirB binding site is required for VirB-dependent modulation of DNA supercoiling of the *PicsP-lacZ* reporter. Analysis of topoisomer distributions of the wild-type *PicsP-lacZ* reporter and its derivative with a mutated VirB binding site (carrying transition mutations at key positions; (47,57)) isolated from *E. coli* in the presence or absence of *virB* induction during growth on an agarose gel containing chloroquine (2.5 μg ml^-1^). Quantification of key lanes (4 & 5, 6 & 7 and 5 & 6) is provided in Figure S6 G-L.

**Figure 3:**
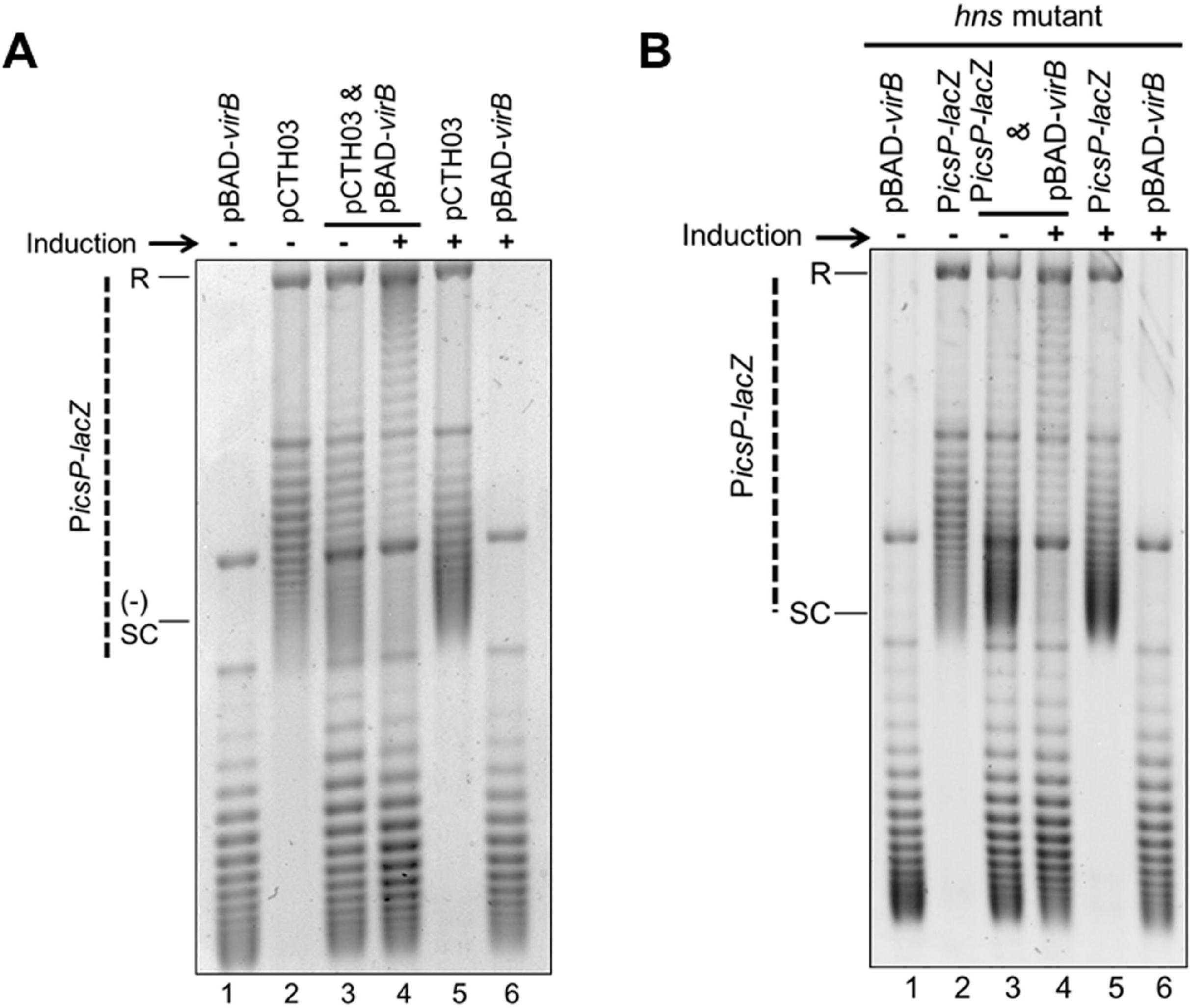
VirB triggers a loss of negative supercoils in *PicsP-lacZ* both in the absence of promoter elements and in an *hns* mutant background. A) Analysis of the topoisomer distribution of *PicsP-lacZ* without promoter elements (pCTH03; (89) in the presence or absence of *virB* induction during growth electrophoresed in the presence of chloroquine (2.5 μg ml^-1^). B) Analysis of the topoisomer distribution of *PicsP-lacZ* reporter isolated from a DH10B *hns* mutant derivative in the presence or absence of *virB* induction during growth electrophoresed in the presence of chloroquine (2.5 μg ml^-1^). Quantification of key lanes (3 & 4) in A and B is provided in Figure S6 S-V.

#### 2D Chloroquine Gel Electrophoresis

Protocols were adapted from Cameron et al., 2011 (80). Initially, DNA samples from *E. coli* were electrophoresed using the same conditions as for 1D gel electrophoresis. Next, the bottom 2 cm of the rectangular gel was removed so that the gel could be rotated 90°, and then the gel was equilibrated to the new chloroquine concentration by soaking in fresh 2x TBE containing 12.5 ug ml^-1^ chloroquine for 4 h in the dark. Subsequently, the DNA was electrophoresed in the dark for 12 h at 2 V cm^-1^. After electrophoresis, the chloroquine was removed from the gel by soaking in fresh 2x TBE for 3 - 4 h, changing the buffer every 30 - 60 min. The gel was then stained with ethidium bromide (100 μg) for 1 h followed by imaging of the gel. For 2D gel electrophoresis of DNA samples isolated from *S. flexneri*, the same electrophoresis conditions were used as for the *E. coli* samples with the following exceptions. In the first dimension, DNA was electrophoresed in the presence of 10 μg ml^-1^ chloroquine for 30 h at 2 V cm^1^, and in the second dimension, 15 μg ml^-1^ of chloroquine for 16 h at 2 V cm^1^ was used. A minimum of two independent experiments were performed, using at least duplicate samples for each experiment. Gels images were captured using a UVP Biospectrum 410 Imaging System and manipulated using Visionworks™ LS Image Acquisition.

### Quantification of gels

Quantification of topoisomer bands was done using AzureSpot Analysis Software version 2.0.062. For the analysis of 1D gels containing samples isolated from cells (Figure S6), the “Analysis Toolbox” was used. Lines were drawn through the centre of lanes starting at the bottom using “draw shapes” feature. Lines were copied using the “duplicate” feature and dragged to another lane so that direct comparisons could be made. The resulting gel images and lane profiles, which indicate the density of each topoisomer band, are shown. For the analysis of 1D gels containing samples from *in vitro* experiments (Figure S8), lanes were identified using “create lanes” feature and boxes were drawn manually with automatic edge detection using “detect bands” feature. Band density (in pixels) was determined after background subtraction using the rolling ball method (with a radius of 200) and the resulting stacked lane profiles are presented. For 2D gel analysis (Figure 4 and Figure S7), the “Analysis Toolbox” was used. Arcs were drawn through the centre of the observed topoisomers using the spline tool in the “draw shapes” feature. Arcs were copied using the “duplicate” feature and dragged to next arc so that direct comparisons could be made. The resulting profiles are shown and indicate the density of each topoisomer band. The ΔLk was calculated by subtracting the topoisomer number with the highest density (defined as the Boltzmann centres) in the VirB^+^ condition (i.e., *virB* inducing conditions in *E. coli* or wild-type *S. flexneri* background) from the topoisomer with the highest pixel density in the VirB^-^ condition (i.e., non-*virB*-inducing conditions in *E. coli*, or *virB* mutant AWY3). The mean and standard deviations were calculated from two independent trials.

**Figure 4.**
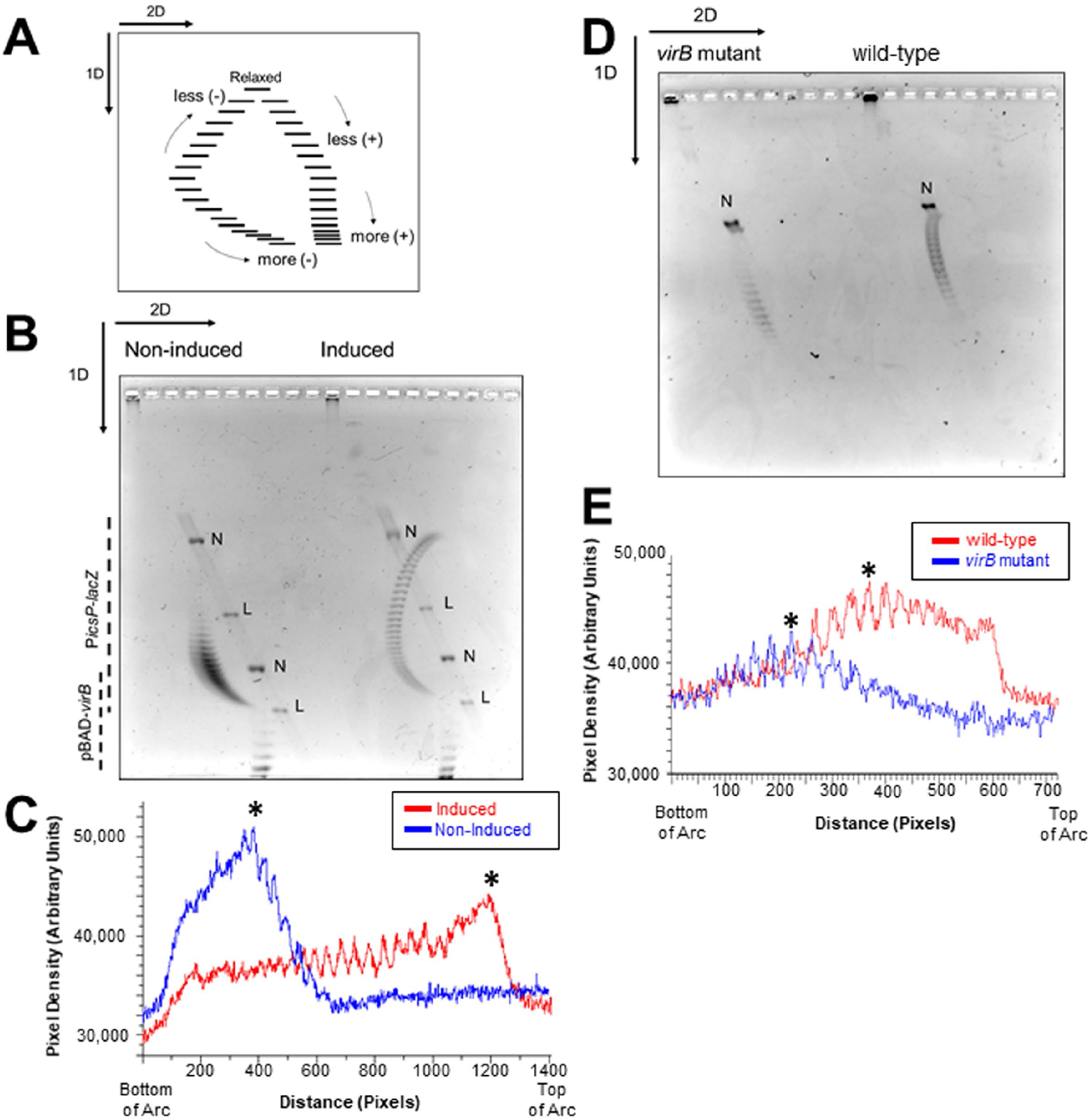
Analysis of *PicsP-lacZ* topoisomer distribution using two-dimensional agarose gel electrophoresis in *E. coli* and *S. flexneri*. A) Schematic of two-dimensional agarose gel electrophoresis used to analyse topoisomer distributions (adapted from (93)). In the first dimension, the fast-migrating topoisomers are more negatively supercoiled while the slow migrating topoisomers are less negatively supercoiled. In the second dimension, the fast-migrating topoisomers are increasingly positively supercoiled. B) Two-dimensional gel electrophoresis of *PicsP-lacZ* reporter isolated from *E. coli* with or without *virB* induction during growth. Two independent trials were performed, representative data are shown. C) Lane trace analysis of topoisomers shown in panel B. Asterisk represents most intense topoisomer in each trace. D) Two-dimensional gel electrophoresis of *PicsP-lacZ* reporter isolated from wildtype *Shigella* or an isogenic *virB* mutant derivative. Two independent trials were performed, representative data are shown. E) Lane trace analysis of topoisomers shown in panel D. Asterisk represents most intense topoisomer in each trace (representative analysis shown) [N=nicked and L=linear]. Arcs used for gel quantification provided in Figure S7.

### *In vitro* VirB supercoiling assays

To analyse the effect of purified VirB-His6 on *PicsP-lacZ* supercoiling *in vitro*, two procedures were used, based on those described in (81–84), with minor modifications. To prepare DNA for *E. coli* topoisomerase I (NEB# M0301) treatment, *PicsP-lacZ* plasmid was isolated from DH10B cells such that the DNA was mostly negatively supercoiled. VirB was incubated with supercoiled plasmid *PicsP-lacZ* in a 1x VirB binding buffer (25 mM HEPES pH 7.6, 100 mM NaCl, 0.1 mM EDTA, 0.5 mM β-mercaptoethanol, 5 % glycerol, 1 mM CTP, 5 mM MgCl_2_). After incubation with VirB, 5 μl was removed from each sample to analyse VirB binding to plasmid DNA using electrophoresis [Figure S4; 0.7% agarose gel in 1x TAE, 100V for 1 h and imaged after staining with ethidium bromide]. Next, 0.2u *E. coli* topoisomerase I was added to the appropriate samples and all samples were incubated for 30 min at 37°C (low amounts of topoisomerase I ensured the retention of some negative supercoils in the plasmid *PicsP-lacZ*). Protein-DNA interactions were disrupted by heating at 65°C for 20 minutes and protein was removed through the addition of 0.8mg ml^-1^ of Proteinase K incubated at 37°C for 1-2h. DNA was then purified using a Promega PCR Clean-Up Kit (Promega #A9282), quantified (Synergy HTX) and analysed by DNA electrophoresis to verify DNA concentrations, before fixed ng of DNA were electrophoresed in the presence of chloroquine (2.5 μg ml^-1^), as described above (1D Chloroquine Gel Electrophoresis; Experimental Procedures). Images were captured (Azure Biosystems, Inc.) after staining with 10 μg ml^-1^ ethidium bromide.

A similar strategy was used to capture VirB-mediated changes in supercoiling using T4 DNA ligase with the following exceptions: i) purified *PicsP-lacZ* was nicked once using Nb.BsrD1 (NEB # R0648S) for 1 h at 65 °C and then purified using phenol/chloroform extraction and ethanol precipitation, ii) nicked DNA was incubated with increasing concentrations of purified VirB-His6 (Monserate Biotech) at 37°C for 30 min in a total reaction volume of 50 μl containing 1x ligase buffer (supplied with T4 DNA ligase; 30mM Tris-HCl pH 7.8, 10mM MgCl_2_, 10mM DTT, 1mM ATP) supplemented with 1mM CTP, iii) after incubation with VirB and sample removal for analysis (Figure S4), the remaining samples were incubated with 6u of T4 DNA Ligase for 2 h at room temperature, and iv) fixed ng of DNA were electrophoresed in the absence of any intercalator. Images were captured (Azure Biosystems, Inc.) after staining with ethidium bromide.

### Promoter activity assays to determine the effect of modulating DNA supercoiling

#### topA mutant assays

MG1655 and its isogenic derivatives DPB923 (*topA* mutant) and DPB924 (*topA-* complemented) (Coli Genetic Stock Center) containing either pBAD18 or pBAD-*virB* (pATM324) and either pMK29 or pAFW04a were grown in triplicate overnight in LB, as described previously. Cells were diluted 1:100 and grown for 3 hours at 37 °C, and then induced with 0.2% (final) L-arabinose for an additional 2 hours. Subsequently, cells were harvested and β-galactosidase activities were determined, as described previously (45).

#### Reporter assays exploiting the twin domain model of transcription

MC4100 or its isogenic *hns* mutant carrying either pAFW04, pParallel, or pDivergent in combination with either pEmpty or pT7-RNAP, were grown in triplicate in LB overnight at 37 °C with aeration (LabLine/ Barnstead MaxQ 4000). Overnight cultures were diluted 1:100 and grown for 2 h at 37 °C, before the addition of a final concentration of 0.002% L-arabinose (85). Cells were then grown for 1 h more before being harvested for both GFP fluorescence and β-galactosidase assays. For the GFP fluorescence, cells were normalised (1.2/OD_600_) and resuspended in 1x PBS. A fixed number of cells were seeded in a black, clear bottomed, 96-well plate (Greiner Bio-One; P/N 655090) and both cell density (OD_600_) and GFP fluorescence (excitation 485, absorbance 528) were measured simultaneously on a Synergy HTX (BioTek) plate reader. β-galactosidase activities were determined as previously described (45).

### VirB protein purification

VirB protein was purified by the Monserate Biotechnology Group (San Diego, CA). Briefly, *E. coli* M15 cells containing pREP4 and pAJH01 plasmids were grown from a glycerol stock in LB broth containing ampicillin (50 μg ml^-1^) and kanamycin (25 μg ml^-1^) until OD_600_ of 0.6 – 0.8. Cells were induced with 1 mM IPTG (IBI Scientific) and grown for 3 hours before harvesting via centrifugation. Cell paste was frozen at −80° C, and then resuspended in 3 ml of lysis buffer per gram of cell paste (20 mM KPO_4_ (pH 7.5), 100 mM NaCl, 1 mM EDTA, 10 mM β-mercaptoethanol, 0.2 mM PMSF). Cells were lysed by two passages through a microfluidizer at 15000 – 20000 psi and the resulting lysate was clarified by centrifugation. NaCl was added to the clarified lysate to a final concentration of 200 mM followed by addition of polyethylenimine (PEI) to a concentration of 0.5 % (v/v). The lysate was stirred for 10 min at room temp and PEI-precipitated nucleic acids were collected by centrifugation. Ammonium sulfate (AS, 35 % w/v) was added to the lysate and precipitated proteins were collected by centrifugation and discarded. A second AS treatment (75 % w/v) of the supernatant was used to precipitate VirB, which was collected by centrifugation. The 75 % AS pellet was resuspended in 80 ml of low salt buffer (LSB, 20 mM KPO_4_ (pH 7.5), 100 mM NaCl, 1 mM EDTA, 1 mM β-mercaptoethanol, 5 % v/v glycerol) and dialyzed against 1 L LSB overnight. Capture of VirB was performed on a 30 mL phosphocellulose (Polysciences, Inc.) column. VirB was eluted using a linear gradient from 0 – 100 % high salt buffer (20 mM KPO_4_ (pH 7.5), 1.5 M NaCl, 1 mM EDTA, 1 mM β-mercaptoethanol, 5 % glycerol) over 7 column volumes. Pooled fractions were diluted 1:1 and purified over a sulfate column (EMD) using the same buffers and gradient as above. Pooled fractions from the sulfate column were dialyzed against LSB before final purification over a TMAE column (EMD). Protein was dialyzed into storage buffer (25 mM HEPES (pH 7.6), 100 mM NaCl, 0.1 mM EDTA, 0.5 mM BME, 5 % glycerol) and flash frozen.

### Statistical Analyses

Statistical calculations were done using IBM^®^ SPSS^®^ Statistics for Windows, version 28.01.0. Analysis of Variance (ANOVA) tests, either one-way, two-way or three-way, were used (noted in figure legends). Throughout this work, statistical significance is represented as *, indicating *p* < 0.05.

## RESULTS

### VirB triggers a loss of negative DNA supercoiling of a *PicsP-lacZ* reporter plasmid *in vivo*

To investigate the ability of VirB to alter DNA topology, the previously described, VirB-dependent *PicsP–lacZ* transcriptional reporter plasmid (45,47) pHJW20, was used (Figure 1A). For this work, the *E. coli* strain DH10B was chosen because i) this strain avoids complications caused by the presence of accessory plasmids in *S. flexneri* cells, and ii) multi-copy plasmids are maintained stably in this strain without risk of recombination (86). Since DH10B does not naturally encode VirB, the *virB* gene was provided *in trans* on an L-arabinose-inducible plasmid, pBAD-*virB*. Initially, the electrophoretic mobility of plasmid DNA isolated from DH10B carrying the *PicsP-lacZ* reporter, pBAD-*virB*, or both in the presence or absence of *virB* inducing conditions was analysed by standard agarose gel electrophoresis. The mobility of single plasmids was not altered by the presence or absence of the inducer (Figure 1B; compare lanes 1 & 6 and lanes 2 & 5), while comparison of samples containing both plasmids to the single plasmid samples revealed that the *PicsP-lacZ* reporter displayed an altered electrophoretic mobility when *virB* was induced (Figure 1B; lane 4). A phenol-chloroform extraction and restriction digest of the samples revealed that the altered mobility of the *PicsP-lacZ* reporter was neither caused by residual VirB protein remaining associated with the reporter (Figure S1A; compare lane 1 with 3 – 5) nor triggered by a VirB-dependent change in size of the *PicsP-lacZ* reporter (Figure S1B; lanes 3 & 4).

Next, the possibility that the observed altered mobility of the *PicsP-lacZ* reporter was caused by a VirB-dependent change in DNA supercoiling was tested. To do this, identically isolated DNA samples were electrophoresed in the presence of chloroquine (2.5 μg ml^-1^), an intercalating agent that resolves plasmid topoisomers by inducing positive supercoiling (78). Chloroquine induces highly negatively supercoiled DNA to become more relaxed, migrating more slowly during electrophoresis; whereas relaxed or positively supercoiled DNA becomes more positively supercoiled and migrates more quickly. In these experiments (Figure 1C), single plasmid controls showed little to no change in their topoisomer distributions, regardless of induction (Figure 1C; compare lanes 1 & 6 and lanes 2 & 5). In contrast, in the DNA samples containing two plasmids, the *PicsP-lacZ* reporter isolated from cells expressing *virB* displayed an altered topoisomer distribution towards the top of the gel when compared to *PicsP-lacZ* isolated from cells without *virB* expression (Figure 1C; compare lanes 3 & 4). These findings suggested that the *PicsP-lacZ* reporter isolated from cells with VirB was more relaxed (less negatively supercoiled) than *PicsP-lacZ* isolated from cells without VirB. This conclusion is supported further by experiments where identical DNA samples were electrophoresed in the presence of a higher concentration of chloroquine (10 μg ml^-1^; Figure S2; quantification provided in Figure S6). Taken together, these data show that the *PicsP-lacZ* reporter has undergone a loss of negative supercoiling when isolated from cells producing VirB.

### VirB-mediated changes in supercoiling are caused by VirB binding to its cognate site

Binding of VirB to its recognition sequence is required for VirB-mediated anti-silencing (46–48,57,58,71). To identify if the interaction of VirB with its target DNA sequences was required for the observed changes in supercoiling, two complementary approaches were used. First, the requirement for the VirB-DNA binding site was examined (Figure 2), and, second, the role of the helix-turn-helix DNA binding domain of VirB was tested.

To identify if the VirB-DNA binding site is required for VirB-mediated changes in DNA supercoiling, the *PicsP-lacZ* reporter derivative bearing a mutated VirB binding site (destroyed by transition mutations), pMIC18 (47), or a wild-type *PicsP-lacZ* control, were analysed using assay conditions similar to those described above. Samples containing both plasmids (Figure 2, lanes 4-7) revealed that only the wild-type P*icsP*-*lacZ* reporter displayed a change in its topoisomer distribution when pBAD-*virB* was induced (Figure 2, compare lanes 4-7), as seen previously (Figure 1C). Notably, mutations within the VirB binding site (pMIC18) in the *PicsP-lacZ* reporter drastically reduced these changes in topoisomer distribution (Figure 2, compare lanes 6 & 7 to lanes 4 & 5, and lanes 5 & 6 to each other; quantification in Figure S6, G-L), demonstrating that the VirB binding site is required for VirB-mediated changes in supercoiling of the *PicsP-lacZ* reporter. Similarly, when VirB K152E, which is compromised in its ability to bind to DNA *in vivo* and *in vitro* (59,71), was expressed in the presence of *PicsP-lacZ* and the topoisomer distribution compared to that generated by wild-type VirB in our assays, the change in topoisomer distribution was much less pronounced (Figure S3A compare lanes 4-7; quantification in Figure S6 M-R), even though equivalent amounts of wild-type VirB and VirB K152E proteins were generated under these assay conditions (Figure S3B). In conclusion, these data demonstrate that the *in vivo* interaction of VirB via its helix-turn-helix binding domain with its cognate DNA-binding site is required for the observed changes in DNA supercoiling.

### VirB-mediated changes in supercoiling are independent of resulting transcription and remodelling of the H-NS:DNA complex

Having demonstrated that VirB-mediated changes in supercoiling are caused by VirB binding to its cognate site *in vivo*, we wanted to establish if this effect was caused directly by the VirB:DNA interaction or if a subsequent VirB-dependent event was responsible. Since transcription is documented to cause localized changes in plasmid DNA supercoiling (87); a process coined the twin-domain model of supercoiling (88), we chose to determine if a VirB-dependent increase in transcription was responsible for the observed changes in supercoiling. To do this, we analysed the topoisomer distribution of a *PicsP-lacZ* reporter, pCTH03, that lacks both VirB-dependent promoters, P1 and P2 (89), but retains the VirB binding site (46,47) (Figure 3A). In DNA samples containing two plasmids (pCTH03 and pBAD-*virB*), the *PicsP-lacZ* derivative, pCTH03, exhibited a change in its topoisomer distribution (Figure 3A, compare lanes 3 & 4) under *virB* inducing conditions, similar to previous results (Figure 1C). Since pCTH03 lacks both characterized VirB-dependent promoters, we concluded that transcription was unlikely to be directly responsible for the observed changes in supercoiling triggered by VirB and that the observed VirB-dependent modulation of DNA supercoiling is likely to precede transcription initiation.

H-NS interactions with DNA are also reported to cause localized changes in DNA topology *in vivo* (90–92). Since VirB functions to relieve transcriptional silencing by H-NS (45, 48,53), likely through the remodelling of H-NS:DNA complexes, we next chose to examine whether the H-NS:DNA complex at the *PicsP-lacZ* reporter was needed for the VirB-dependent changes in topoisomer distribution observed in our assays. To address this, the topoisomer distributions of full-length *PicsP-lacZ* reporter and pBAD-*virB* were analysed when isolated from an *E. coli* DH10B *hns* mutant derivative. In DNA samples containing both the *PicsP-lacZ* reporter and pBAD-*virB* plasmids isolated from the DH10B *hns* mutant, the reporter plasmid, *PicsP-lacZ*, again displayed an altered topoisomer distribution (Figure 3B, compare lanes 3 & 4) but only under inducing conditions. These results allowed us to conclude that the observed VirB-dependent alteration in topoisomer distribution of *PicsP-lacZ* occurs even in the absence of functional HNS. Thus, the VirB-dependent change in topoisomer distribution of *PicsP-lacZ* that occurs *in vivo* (Figure 1C, & Figure 2) relies on specific VirB:DNA interactions and precedes any VirB-dependent remodelling of the H-NS:DNA complex and the resulting transcription from the *icsP* promoter.

### A loss of negative *PicsP-lacZ* supercoiling occurs in *S. flexneri* with native VirB levels

Having demonstrated that the ectopic expression of plasmid-borne inducible *virB*, led to changes in changes in DNA supercoiling of the *PicsP-lacZ* reporter, we next investigated if native levels of VirB were sufficient to cause detectable changes in supercoiling. Although early attempts to analyse DNA isolated from *S. flexneri* using standard, one-dimension gel electrophoresis were hindered by the presence of accessory plasmids and poor quality of the isolated DNA (data not shown), the use of two-dimensional DNA gel electrophoresis resolved these problems and allowed high resolution of negatively and positively supercoiled topoisomers within each DNA sample. Briefly, in the first dimension of electrophoresis, the fast-migrating topoisomers could be either positively or negatively supercoiled while the slower migrating topoisomers would be more relaxed. In the second dimension, with a higher concentration of chloroquine, all topoisomers regardless of whether they were originally positively or negatively supercoiled become more positively supercoiled due to the presence of the intercalator, thus allowing them to be resolved (Figure 4A; adapted from (93)).

Prior to testing DNA samples isolated from *S. flexneri*, we first chose to analyse DNA samples containing both the *PicsP-lacZ* reporter and pBAD-*virB* isolated from *E. coli* DH10B (complexes like those analysed in Figure 1C & S2) using two-dimensional gel electrophoresis. These proof of principle experiments would allow direct comparison with DNA samples subsequently isolated from *S. flexneri*. In the absence of *virB* induction, the *PicsP-lacZ* reporter topoisomers were all negatively supercoiled (Figure 4B) and arranged in a tight distribution (Figure 4C). However, in the presence of *virB* induction, the *PicsP-lacZ* topoisomers were comparatively more relaxed (Figure 4B) and spread in a wider distribution (Figure 4C). The average change in linking number (ΔLk) from the non-inducing to *virB* inducing conditions is ΔLk = +21 ± 1.4, indicating that VirB triggers a loss of negative supercoiling consistent with our previous assays (Figures 1C and S2).

Next, we analysed the change in topoisomer distribution of the *PicsP-lacZ* reporter isolated from wildtype *S. flexneri* and an isogenic *virB* mutant using the new DNA isolation protocol and two-dimension gel electrophoresis. The *PicsP-lacZ* reporter isolated from the *virB* mutant were all negatively supercoiled (Figure 4D) and displayed a tight distribution of *PicsP-lacZ* topoisomers (Figures 4E). In contrast, the *PicsP-lacZ* reporter isolated from wild-type *S. flexneri* cells displayed a relatively broad distribution of relaxed topoisomers (Figure 4D & E) with an average ΔLk = +4.0 ± 0. These data demonstrate that native levels of VirB in *S. flexneri* are indeed sufficient to trigger a loss of negative supercoiling of the *PicsP-lacZ* reporter similar to our initial observations made in *E. coli*, albeit, to a lesser degree.

### Topoisomerase I is required to capture VirB-dependent changes in supercoiling *in vivo*, but antisilencing proceeds in its absence

All experiments, thus far, indicated that plasmids bearing a VirB DNA-binding site undergo a loss of negative supercoils in the presence of wild-type VirB when recovered from bacterial cells (Figures 1B & C, 2, S2, 3, 4B & D). But if VirB was directly causing the change in DNA supercoiling, the VirB-induced changes would be lost when the protein was removed during plasmid preparation. Thus, either VirB was acting like a topoisomerase, which seemed unlikely based on lack of homology, or a cellular topoisomerase was “fixing” the transient topological changes caused by VirB, so that the changes could be detected in isolated plasmids. Since the topological change observed in our assays was a loss of negative supercoils, it seemed likely that topoisomerase I might be involved.

To test this, we exploited a wild-type *E. coli* strain (MG1655) or a mutant derivative (DPB923) lacking topoisomerase I (the *topA10* mutant) (94). Wild-type *PicsP-lacZ* plasmid and the inducible pBAD-*virB* were introduced into each of these backgrounds and the plasmids were analysed as described previously using 1D agarose gel electrophoresis in the presence of chloroquine. Under these conditions, the induction of VirB in the wild-type strain caused a change in the topoisomer distribution of *PicsP-lacZ* like that seen before (although, the quality of the DNA isolated from this *E. coli* strain was lower than that isolated from DH10B, as expected; Figure 5A). In contrast, in the *topA10* mutant background these pronounced topological changes were absent. Instead, the most striking changes were seen in pBAD-*virB* under conditions of induction (likely caused by transcription from pBAD). Thus, these data strongly suggest that topoisomerase I is responsible for capturing the VirB-dependent changes in DNA supercoiling observed in plasmids isolated from cells.

**Figure 5.**
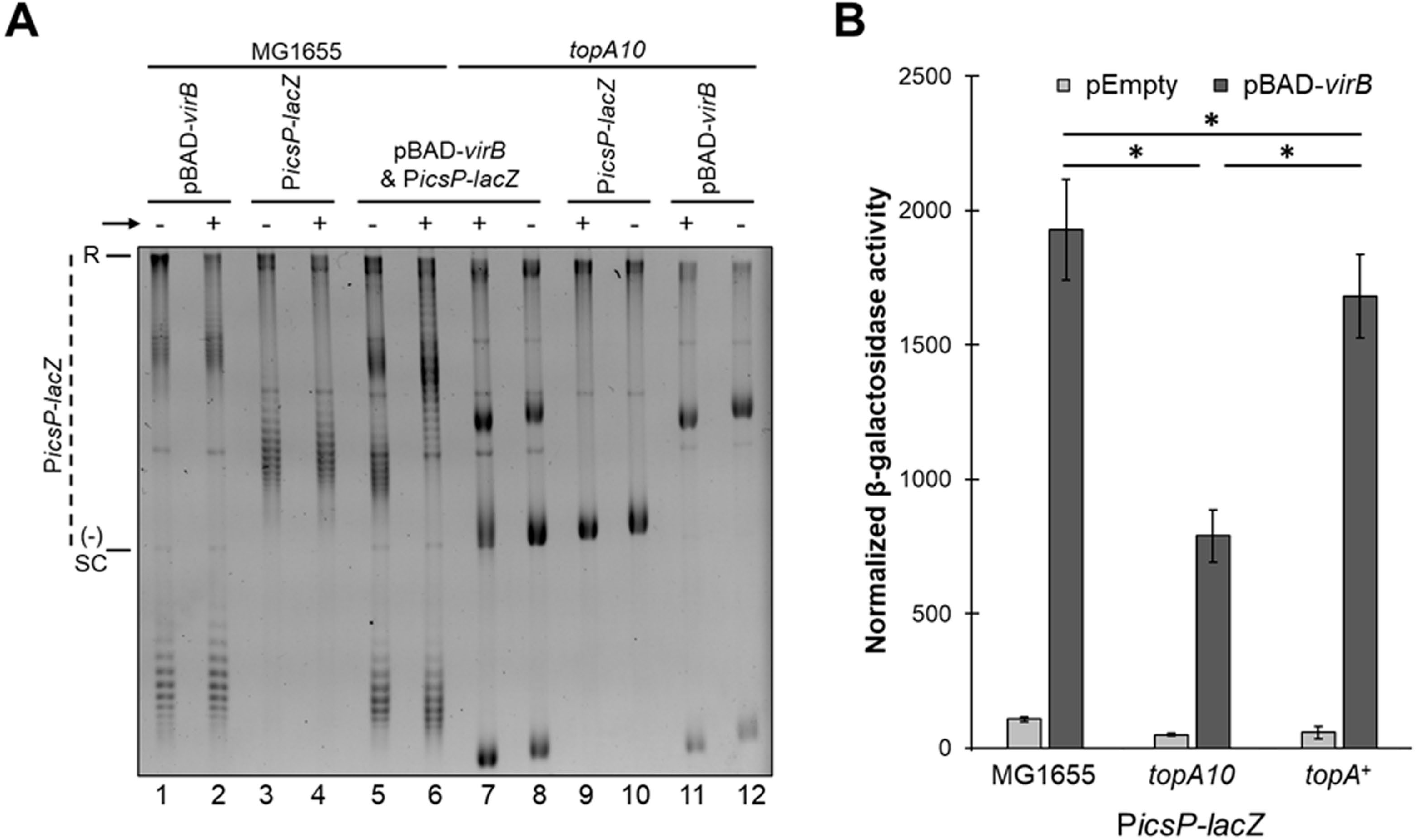
Topoisomerase I captures VirB-dependent changes in supercoiling *in vivo* but is not needed for anti-silencing. A) Analysis of the topoisomer distribution of *PicsP-lacZ* in *E. coli* MG1655 and the *topA10* mutant derivative in the presence or absence of *virB* induction during growth electrophoresed on 1% agarose in the presence of chloroquine (2.5 μg ml^-1^). Lanes 1-4 and 9-12 contain 125 ng of DNA, whereas lanes 5-8 contain 250 ng. Quantification of key lanes (5–8) on gel shown in A is provide in Figure S6 W-Z. B) Normalised promoter activity of *PicsP-lacZ* measured in either wild-type *E. coli, topA10* mutant (12), or *topA* complemented strain (12). Assays were done with three biological replicates and repeated three times. Representative data are shown. Significance was calculated using a two-way ANOVA, with post-hoc Bonferroni, *, *p*<0.05. Complete statistical analysis provided in Table S3.

Having identified that topoisomerase I was involved, the next step was to examine the role that topoisomerase I plays in transcriptional anti-silencing by VirB. The activity of the well-characterized VirB-dependent promoter *PicsP* (46,47) was measured using our plasmid reporter *PicsP-lacZ* in the wild type *E. coli* strain (MG1655), the *topA10* mutant (DPB923) and a complemented strain (DPB924). In the topoisomerase mutant, *PicsP* activity in the absence of VirB was lower than in wild-type (1.3 ± 0.1-fold). This suggests that the accrual of negative supercoils in the absence of topoisomerase I, modestly downregulates the activity of *PicsP*. Furthermore, the absence of topoisomerase I also decreased in VirB-dependent activity of *PicsP* (1.8 ± 0.2 fold lower), but importantly, VirB-dependent regulation was still observed in all strains tested (Figure 5B). Thus, topoisomerase I is not an absolute requirement for VirB-dependent regulation, although it may contribute to the regulatory effect of VirB *in vivo*.

### In vitro, VirB induces positive supercoils in plasmid DNA

Next, we chose to examine whether VirB could trigger a change in the topoisomer distribution of plasmids *in vitro*. Initial experiments revealed that if a purified VirB stock capable of binding to its cognate site was incubated with *PicsP-lacZ* for 30 mins at 37°C, after the VirB protein was removed, no VirB-dependent change in the topoisomer distribution of *PicsP-lacZ,was* observed by 1D chloroquine gel electrophoresis (data not shown). This supported our theory that VirB does not possess topoisomerase activity. Moreover, these findings indicated that we would likely need a way to fix the VirB-induced change in DNA supercoiling if we were to characterize the VirB mediated effect *in vitro*.

To do this, two complementary approaches were used, employing either topoisomerase I or T4 DNA ligase as the “fixing agent” (Figure 6; corresponding EMSAs in Figure S4). In the first approach, negatively supercoiled plasmid DNA was incubated with VirB and then exposed to low concentrations of topoisomerase I (Figure 6A). Low levels of topoisomerase I would mildly relax the plasmid DNA and allow us to determine the torsion effect that VirB was having on the DNA. As expected, negatively supercoiled DNA incubated without VirB or topoisomerase I, ran at the bottom of the gel (lane 1). When negative supercoiled DNA was incubated without VirB but with the low concentration of topoisomerase I, the topoisomer distribution shifted up the gel, due to some negative supercoils being removed from the plasmid by the topoisomerase (lane 2). The addition of increasing concentrations of VirB followed by topoisomerase treatment, enhanced this effect: topoisomers were initially seen to shift up the gel with increasing concentration of VirB (lanes 3 & 4), but at the highest concentration of VirB (lane 5) this effect was reversed, as the DNA started to accrue positive supercoils (quantification provided in Figure S8A). The altered distribution of topoisomers with increasing concentrations of VirB is wholly consistent with VirB introducing positive supercoils into the *PicsP-lacZ* plasmid.

**Figure 6.**
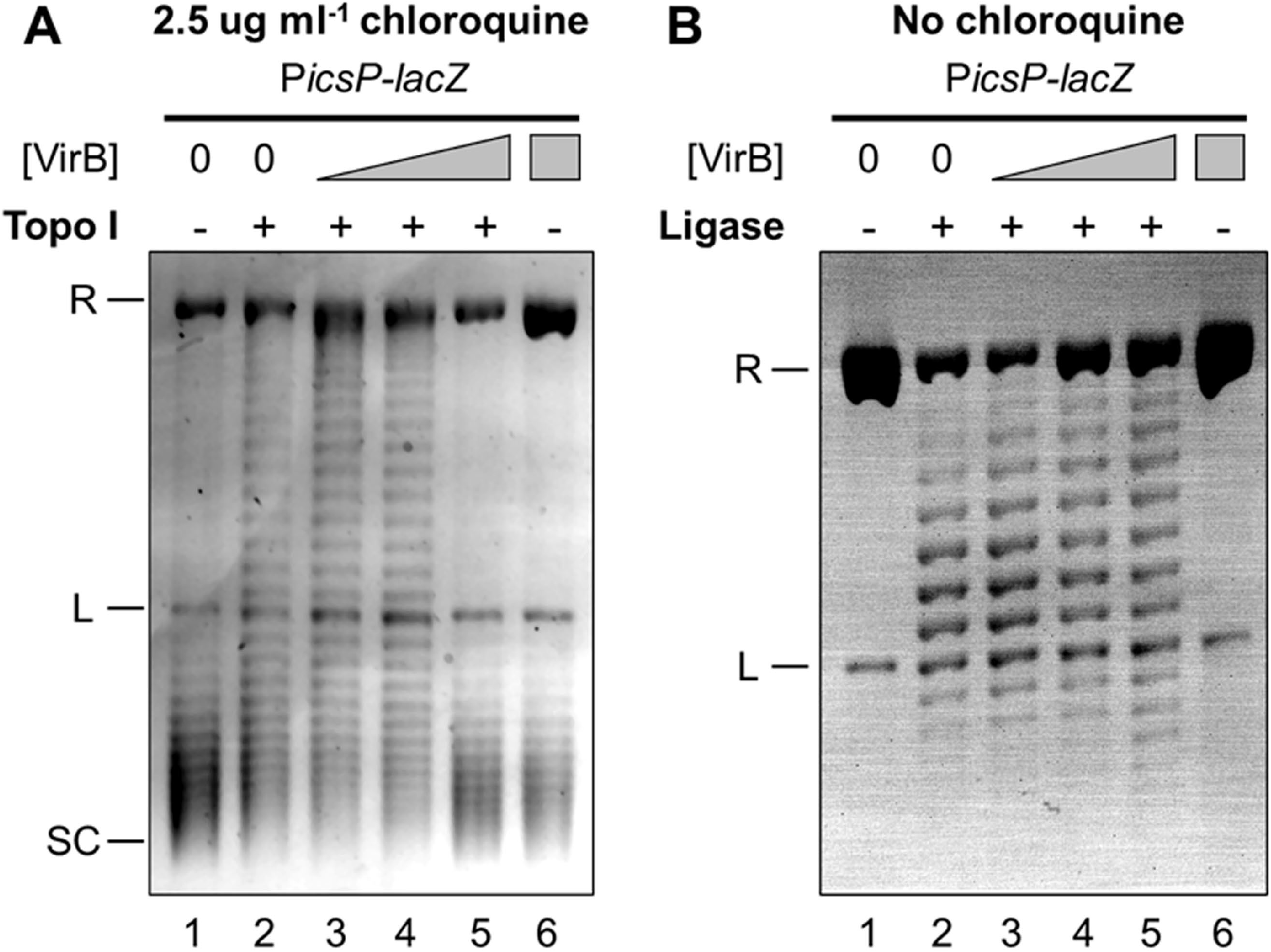
VirB introduces positive supercoils into plasmid DNA *in vitro*, captured by two complementary approaches. A) VirB-dependent changes on a negatively supercoiled *PicsP-lacZ* plasmid captured using *E. coli* topoisomerase I. Increasing concentrations of VirB (lanes 1 and 2 = 0 μM, lane 3 = 0.12 μM, lane 4 = 0.35 μM, lanes 5 and 6 = 0.59 μM) were incubated with negatively supercoiled DNA (3.3 nM) and then samples were mildly relaxed with topoisomerase I (0.2 units). Deproteinised DNA (150 ng) was loaded on an agarose gel containing chloroquine. Quantification of lanes is given in Figure S8 A & B. B) VirB-dependent changes on nicked P*icsP-lacZ* plasmid captured using T4 DNA Ligase. Nicked DNA (3.0 nM) was incubated with increasing concentrations of VirB (lanes 1 and 2= 0 μM, lane 3 = 0.12 μM, lane 4 = 0.24 μM, lanes 5 and 6 = 0.35 μM) and then treated with DNA ligase (6 units). Deproteinised DNA (150_ng) was loaded on an agarose gel with no chloroquine. R = relaxed, L = linear, SC = supercoiled. Representative images of three trials are shown. Quantification of lanes is provided in Figure S8 C & D.

In the second approach, nicked and hence relaxed plasmid DNA (nicked with Nb.BsrDI at one site located ~2.2 kb away from the VirB-binding site) was incubated with VirB and then treated with DNA ligase (Figure 6B). Resulting topoisomers were then separated on a 1D agarose gel without any intercalator. The absence of any intercalator would ensure that any positive supercoils introduced into the covalently closed circular DNA by VirB could be detected. Nicked plasmid incubated without VirB or ligase ran as relaxed DNA at the top of the gel (lane 1). When nicked plasmid was incubated without VirB but with DNA ligase, the plasmid DNA became a covalently closed circle with some supercoils (lane 2; a similar effect was observed in (83)). When ligase treatment followed incubation with the lowest concentration of VirB, the distribution of topoisomers in the covalently closed product moved up the gel (lane 3) but began to move down the gel with the second highest concentration (lane 4) and further down the gel with the highest concentration (lane 5) (quantification provided in Figure S8B). While these changes in the distribution of topoisomers were visible in this assay, clearly these results were somewhat hampered by the increasing concentrations of VirB inhibiting the “fixative” in this assay – the DNA ligation reaction (an increasing population of relaxed/nicked DNA was seen at the top of the gel with increasing concentrations of VirB after ligase treatment). Regardless, these findings are again consistent with VirB introducing positive supercoils into the *PicsP-lacZ* plasmid.

### Loss of negative supercoils is sufficient to relieve H-NS mediated transcriptional silencing

Since, VirB introduces positive supercoils into *PicsP-lacZ* through engagement of its DNA binding site, we next chose to examine if the introduction of positive supercoils would relieve H-NS-mediated silencing of the *icsP* promoter. To test this, we exploited the twin-domain model of transcription (87,88) to modulate local changes in DNA supercoiling at the *icsP* promoter. Two plasmid reporter constructs were built (Figure 7A), each was engineered to carry a *gfp* reporter under the control of a highly active T7 polymerase-dependent promoter (Pϕ10). This genetic locus was then inserted upstream of the *icsP* promoter in the *PicsP-lacZ* reporter in either a divergent or parallel orientation with respect to the *lacZ* gene (pDivergent and pParallel, respectively). Additionally, five consecutive *rrnB* T1 transcriptional terminators were positioned downstream of the *gfp* gene in each case to prevent transcriptional read-through. Based on the twin domain model, transcription from Pϕ10 would create positive supercoils ahead and negative supercoils behind the elongating T7 polymerase. Thus, we could modulate DNA supercoiling locally, within the *icsP* promoter region, in a VirB-independent manner, simply by inducing the expression of *gene1*, which encodes T7 polymerase.

**Figure 7.**
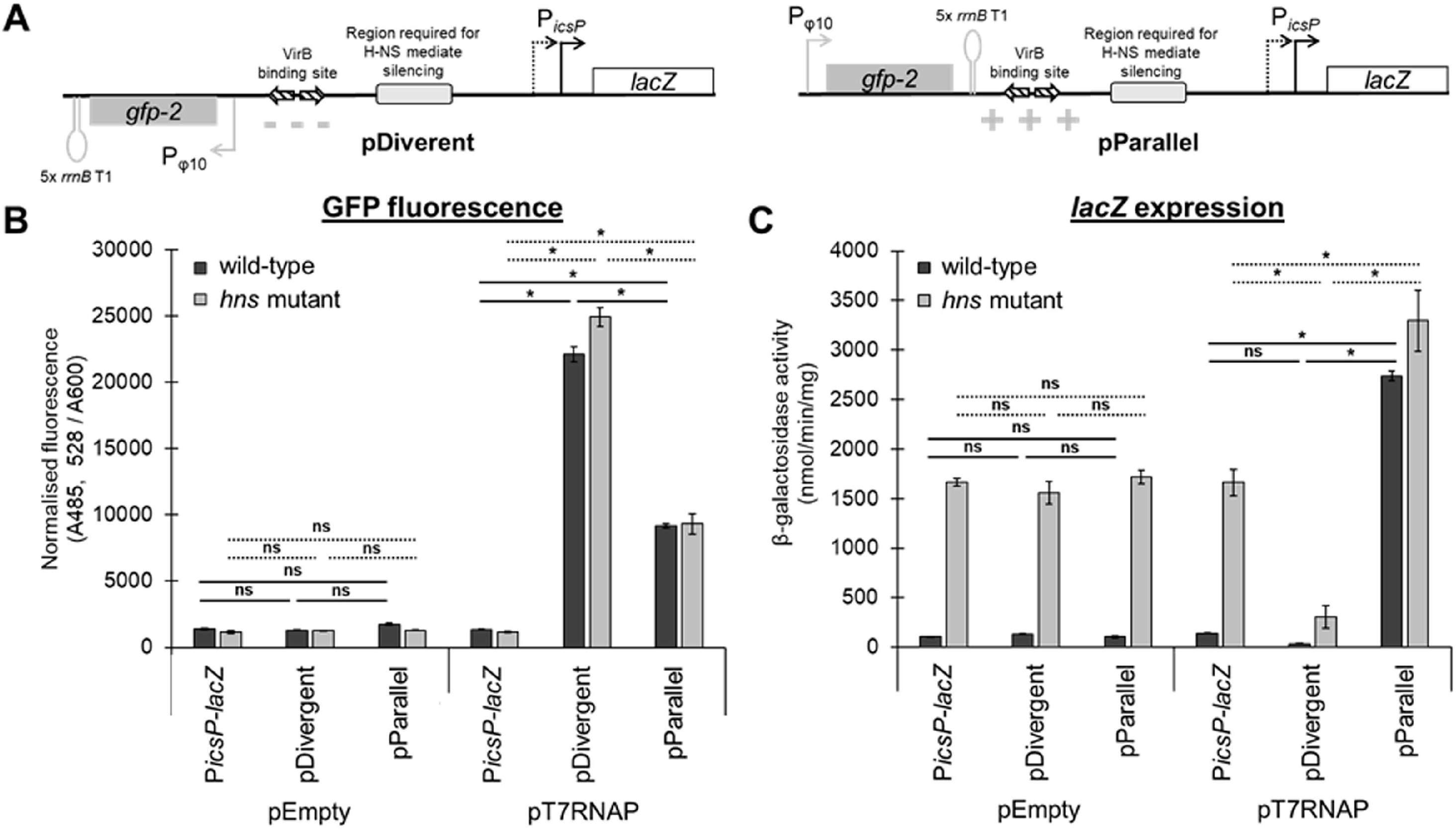
H-NS-mediated silencing is either alleviated or enhanced by an active T7-dependent promoter when positioned upstream of *PicsP* in two orientations, independently of VirB. A) Schematic of pDivergent and pParallel T7 RNAP-dependent Pφ10 cloned in two orientations upstream of *PicsP-lacZ*. The localized changes in supercoiling generated in the *icsP* promoter region predicted by the twin-domain model of transcription-coupled supercoiling (87) are depicted for each construct. B) Normalised GFP fluorescence is depicted as a proxy for Pφ10 activity for each reporter construct (*PicsP-lacZ*, pDivergent, and pParallel) with either pEmpty or pT7-RNAP in *E. coli* wild-type or *hns* mutant. C) β-galactosidase activity as a proxy for *icsP* promoter activity for each of the reporter constructs (*PicsP-lacZ*, pDivergent, or pParallel) either with pEmpty or pT7-RNAP in *E. coli* wild-type or *hns* mutant. Assays were done with three biological replicates and repeated three times. Representative data are shown. Significance was calculated using a three-way ANOVA with post-hoc Bonferron, *, p<0.05 (solid lines compare wild-type and dashed lines compare *hns* mutant). Complete statistical analysis provided in Table S4.

As an initial step, we examined GFP fluorescence as a proxy for ϕ10 promoter activity in each construct (Figure 7B). As expected, average GFP fluorescence was significantly higher in the presence of T7 polymerase than in its absence. With T7 polymerase, GFP fluorescence was not more than 2.5-fold different, regardless of the orientation of the *gfp* gene or the cellular background. Next, we examined β-galactosidase activity as a proxy for *PicsP* activity (Figure 7C). Strikingly, when T7 polymerase was induced, the activity of *PicsP-lacZ* on pParallel increased 19.4 ± 0.6 fold in cells with H-NS (dark bars) but only 2.0 ± 0.3 fold in the *hns* mutant (light bars). In contrast, the activity of *PicsP-lacZ* on pDivergent dropped by 4-6 fold from baseline, regardless of the cell background (Figure 7C). Taken together, these data reveal the sensitivity of *PicsP-lacZ* to changes in DNA supercoiling but, more importantly, reveal that H-NS-mediated silencing can clearly be relieved by the introduction of positive supercoils into the promoter region, whereas negative supercoils seem to hinder promoter activity (consistent with data shown in Fig. 5B), irrespective of whether H-NS is present or not.

To further assess the impact that the introduction of positive supercoils has on the activity of *PicsP*, we also examined *PicsP* activity in a *Shigella virB* mutant in the presence of the DNA gyrase inhibitor, novobiocin (Figure S5). Poisoning of DNA gyrase with novobiocin would be predicted to lead to a loss of negative supercoils from the DNA, mirroring the effect that VirB has on DNA (Figures 1C, 2, 3, 4B, 4D & S2). Indeed, in our experiments utilizing novobiocin, normalised *PicsP* activity increased with the addition of novobiocin. Thus, these data, along with those shown in Figure 7C, are consistent with the introduction of positive supercoils relieving H-NS mediated silencing of our reporter *PicsP-lacZ*.

In sum, these studies clearly demonstrate that a loss of negative supercoiling relieves H-NS mediated silencing of the *icsP* promoter and that the anti-silencer VirB triggers similar topological effects when it binds to its specific DNA site on the *PicsP-lacZ* reporter (Figures 1C, 4B, 4D & S2). Thus, we posit that the modulation of DNA supercoiling triggered by VirB is a critical mechanistic component of its ability to act as a transcriptional anti-silencer. A model, based on this work that provides our current understanding of transcriptional anti-silencing by VirB at *PicsP*, is presented (Figure 8) and discussed below.

**Figure 8.**
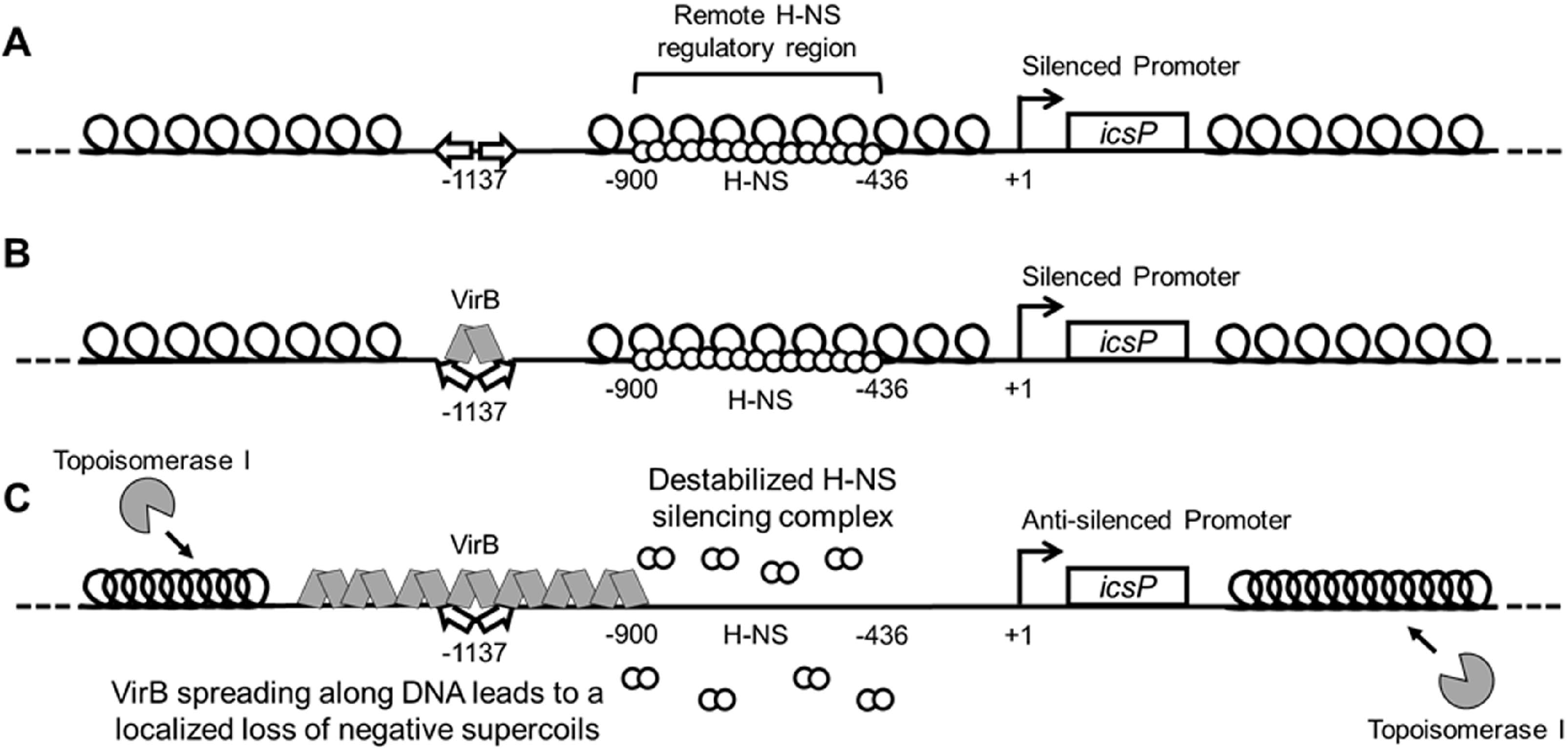
Model showing the proposed mechanism of VirB-induced supercoiling and its alleviation of H-NS-mediated transcriptional silencing. A) At ambient temperatures, 30°C, the *PicsP-lacZ* reporter is negatively supercoiled and transcriptionally silenced by H-NS, which binds to a remote region located between 900 to −436 relative to the primary *icsP* TSS (46). B) Upon a switch to 37°C, VirB docks to its binding site as a dimer (46, 47,59), inducing a slight bend in the DNA (58). C) VirB subsequently oligomerizes along DNA bi-directionally (46, 59,71). VirB spreading along the DNA toward H-NS is required for transcriptional anti-silencing (46), leading to a loss of negative DNA supercoils in the region bound by H-NS. This loss of negative supercoils destabilizes or remodels the H-NS-silencing complex, rendering the promoter permissive for transcription. The topological changes mediated by VirB, lead to the accrual of negative supercoils in unoccupied regions of the plasmid, which attract topoisomerase I to restore negative supercoiling to normal levels (fortuitously, this allowed VirB-mediated changes in supercoiling to be captured in plasmid DNA in our assays). Thus, the anti-silencing activity of VirB is mediated directly through VirB:DNA interactions, topoisomerase I may contribute, but is not the primary driver of this regulatory process.

## DISCUSSION

Mechanisms that counter transcriptional silencing by nucleoid structuring proteins, like H-NS and its functional homologs, are poorly understood. Yet these regulatory events are vital for broad swaths of bacterial life because they control bacterial metabolism and physiology at the genetic level. Anti-silencing proteins function to either liberate or remodel sequestered DNA so that silenced DNA can become transcriptionally active. In this work, we probe the molecular mechanism that underpins the transcriptional anti-silencing activity of VirB, a critical controller in the transcriptional cascade that controls *Shigella* pathogenicity and virulence (50,95). Even though bacterial anti-silencing proteins are diverse, it seems probable that the mechanistic underpinnings of countering NSPs are shared by anti-silencing proteins. Thus, the goal of this study was two-fold: to provide new insight into the molecular mechanism that controls virulence gene expression in the important human pathogen *Shigella* and, more broadly, to enhance our mechanistic understanding of transcriptional anti-silencing, a key bacterial process (13).

In this work, we show that the *Shigella* anti-silencing protein VirB mediates a loss of negative DNA supercoiling from the plasmid-borne, VirB-regulated *PicsP-lacZ* reporter *in vivo*. These changes are not caused by a VirB-dependent increase in transcription, nor do they require the presence of H-NS. Instead, they depend upon the interaction of VirB with its DNA binding site (57); a necessary first step in VirB-dependent gene regulation (59). *In vivo*, VirB-dependent changes in DNA supercoiling are fixed into plasmids bearing its recognition site by topoisomerase I. But topoisomerase I is not a requirement for VirB-dependent regulation, because VirB-dependent regulation is still observed in a *topA10* mutant strain. Instead, VirB:DNA interactions are responsible for generating positive supercoils transiently in plasmid DNA, events which are captured in this work using two complementary *in vitro* approaches. To further examine the significance of the topological changes mediated by VirB, the impact that modulation of DNA supercoiling has on H-NS-mediated transcriptional silencing was investigated. By exploiting the twin-domain model of transcription (88), a localized loss of negative supercoils generated in the region of *PicsP* predicted to be bound by H-NS, was found to be sufficient to alleviate H-NS-mediated transcriptional silencing, even in the absence of VirB. Furthermore, the addition of novobiocin, which poisons DNA gyrase and leads to the accrual of positive supercoils, also alleviated H-NS-mediated silencing of *PicsP*, in a VirB-independent manner. To the best of our knowledge, this is the first time that H-NS-mediated silencing of transcription has been shown to be relieved by a loss of negative supercoils in the region bound by H-NS. Additionally, our work demonstrates that the introduction of positive supercoils by DNA binding proteins like VirB, may provide a mechanism to overcome the repressive effects of H-NS, a widely distributed nucleoid structuring protein in bacteria.

Our model of transcriptional anti-silencing by the VirB protein (Figure 8), relies on the data presented in this study, along with previous research from our team and others (45–48,71). *In vivo*, at ambient temperatures, the *icsP* promoter is negatively supercoiled and transcriptionally silenced by H-NS (Figure 8A) (45). The region required for H-NS mediated silencing lies between −900 and −436 with respect to the primary transcription start site (+1) (46,89). Upon a switch to 37°C, VirB is produced (50). Under these conditions, VirB recognizes and binds to its cognate site (57) centred at −1137 (46,47), inducing a slight bend in the DNA (58) (Figure 8B). From the work presented here, the binding of VirB to its site is essential for the subsequent change in DNA supercoiling, as site directed mutagenesis of the VirB-DNA recognition site or use of the VirB K152E DNA-binding mutant significantly reduces/inhibits these changes (Figures 2 & S3). Importantly, we observe that the resulting loss of negative supercoils occurs at physiological levels of VirB (Figures 4D & E), which agrees with our observations made when *virB* is overexpressed. The mechanistic steps, however, between VirB binding DNA, and the net loss of negative supercoils that ultimately relieves H-NS mediated silencing, remain less clear.

Insight into the mechanistic details that link VirB binding to DNA with the net loss of negative supercoils is provided by our prior research on VirB and the relationship of VirB with members of the ParB superfamily. VirB forms higher order oligomers both *in vitro* and *in vivo* (58,59). Thus, while the DNA binding activity of VirB is essential for transcriptional anti-silencing of virulence plasmid genes (46, 57–59,71), VirB oligomerization or spreading on DNA is likely a critical component of its anti-silencing activity. Evidence for VirB oligomerization or spreading has been observed by DNase I protection assays where VirB protects long swaths of DNA flanking its DNA recognition site (46). Additionally, VirB is a member of of the ParB superfamily (48,96). These proteins function in the faithful segregation of newly replicated DNA to daughter cells, during cell division. During this process, ParB spreads bi-directionally along DNA (63,64) in a process recently demonstrated to require a CTP ligand (69, 70,97). Notably, the spreading activity of P1 phage ParB (63) and the related SopB of F plasmid, both trigger a loss of negative supercoiling of plasmids bearing their respective DNA binding sites (65–67) and a protein docked upstream and downstream of the ParB binding site prevents these supercoiling changes (65,66).

Considering the parallels between VirB and members of the ParB superfamily, the spreading of VirB along DNA is clearly implicated in the modulation of DNA supercoiling and so it features in our model (Figure 6C). Whether spreading of VirB directly or indirectly triggers these topological changes by recruiting topoisomerases to the DNA, up to this point has been less clear.

In this study, we consider the role that topoisomerases play in transcriptional anti-silencing by VirB and VirB-mediated topological changes triggered through its engagement of DNA. VirB:DNA interactions can be seen to generate positive supercoils in DNA both *in vivo* and *in vitro*, although these changes are transient in nature because they are only detected when a “locking agent” is present. *In vivo*, topoisomerase I serves as this locking agent because changes in DNA supercoiling are not observed in plasmids with VirB binding sites if they are isolated from VirB containing cells lacking topoisomerase I (Figure 5A). *In vitro*, either topoisomerase I or DNA ligase can serve as this locking agent (Figure 6 & Figure S8). The capturing of VirB-mediated supercoils by topoisomerase I *in vivo* raised the possibility that this enzyme was not just a fixer of the topological change mediated by VirB, but directly responsible for the removal of negative supercoils required to overcome H-NS-mediated silencing. This idea was refuted, however, by promoter activity assays in a *topA* mutant, where VirB-dependent regulation of *PicsP* was still observed in cells lacking topoisomerase I, albeit to a lesser degree (Figure 5B). Although topoisomerase III or topoisomerase IV are present in the *topA* mutant, and may functionally substitute for topoisomerase I in its absence (98), VirB-mediated changes in DNA supercoiling were not captured in the absence of topoisomerase I *in vivo* (Figure 5A). Thus, it seems most likely that topoisomerase I functions alone and only as a consequence of the changes in DNA topology that are directly caused through VirB:DNA interactions. Thus, we conclude that topoisomerase I is not the primary driver of the topological changes that relieves H-NS mediated silencing. Instead, the VirB-mediated localized stiffening of the DNA as it spreads along the DNA helix into a region occupied by H-NS is most likely required (Figure 8).

Support for this, comes from previous experiments where we introduced a protein roadblock on either side of the VirB binding site in our *PicsP-lacZ* reporter. When the roadblock was positioned between the VirB binding site and the region required for H-NS mediated silencing, VirB-dependent regulation of the *icsP* promoter was blocked (46). Importantly, this blockage was not observed if the LacI protein was docked upstream of the VirB binding site (46). These data suggest that while spreading and subsequent localized loss of negative supercoils by VirB occurs in both directions, only when the loss of negative supercoils occurs in the region bound by H-NS is the anti-silencing activity of the VirB protein revealed. In summary, our findings are consistent with a model (Figure 8) where VirB bidirectionally spreads along DNA, which directly causes a stiffening of the DNA filament such that negative supercoils are lost in the immediately adjacent DNA. If H-NS is in the path of these topological changes then H-NS-mediated silencing is lost. Secondarily, the spreading of VirB and concomitant loss of negative supercoils in one region of DNA, ultimately squeezes negative supercoils into other unbound regions of the plasmid. It is this accrual of negative supercoils that likely attracts DNA topoisomerase I so that the superhelical density of the plasmid in the presence of VirB can be restored to normal levels.

To conclude, in this study we have demonstrated that native levels of VirB trigger a loss of negative supercoiling when VirB binds to a plasmid bearing its recognition site *in vivo or in vitro* (Figures 1B & C, 2, S2, 3, 4B & D). We further show that a loss of negative supercoiling, independent of VirB, can alleviate transcriptional silencing mediated by H-NS (Figures 5 & 6). Our work points to a VirB-dependent loss of negative supercoils being the direct mechanism that relieves H-NS mediated silencing of virulence genes mediated in *Shigella*. While we have focused on VirB in this work, our findings have implications for others studying transcriptional anti-silencing, because although anti-silencers are a diverse group of DNA binding proteins (examples include Ler, LeuO, RovA, SlyA, VirB and VirF) the modulation of DNA supercoiling may be a common mechanistic underpinning of transcriptional anti-silencing. The interplay between transcriptional anti-silencers, nucleoid associated proteins, and DNA topology, lies at the heart of bacterial pathogenesis and physiology and controls the ability of bacteria to respond to environmental changes (3,4,25,100). Future studies focused on gaining a comprehensive understanding of the activities of transcriptional anti-silencers, nucleoid associated proteins, and how they affect DNA topology are clearly needed. We also predict that the molecular underpinnings of transcriptional anti-silencing may be shared, and that common themes will emerge as these important regulatory processes are further studied in *Shigella*, and other bacteria.

## SUPPLEMENTARY DATA

Supplementary Data are available at NAR online.

## DATA AVAILABILITY

The authors confirm that the data supporting the findings of this study are available within the article and its supplementary materials.

## ACKNOWLEDGEMENTS

We thank Monserate Biotechnology Group for purification of VirB-His6, Dr. N. Pat Higgins for guidance with choloroquine gels, Dr. Justin Courcelle for advice on the quantification of topoisomers and Dr. Boo Shan Tseng for advice on statistical analyses. We thank lab members past and present for insightful discussions on the work presented in this manuscript and the four anonymous reviewers for their ideas, comments and helpful suggestions.

## FUNDING

This work was supported by the National Institute of Health [R15 AI090573 to H.J.W, the UNLV Genomics Core Facility, used throughout this study, & J.C.D., were supported by P20 RR-016464 from INBRE Program of the National Center for Research Resources]. The content of this paper is solely the responsibility of the authors and does not necessarily represent the official views of NIH. M.A.P. & J.A.M. were recipients of a Higher Education Graduate Research Opportunity Fellowship from the Nevada Space Grant Consortium NASA Training Grant NNX15AI02H and several fellowships and grants from UNLV and affiliated associations like the Association of Biology Graduate Students and the Graduate & Professional Student Association. T.M.G. was a recipient of a U.S. Department of Education GAANN fellowship (P200A210055). These funding sources had no role in the study design, data collection and interpretation, or the decision to submit the work for publication. Funding for open access charge: National Institutes of Health R15 AI090573.

## CONFLICT OF INTEREST

There are no conflicts of interest to declare.

## Notes

### Competing Interest Statement

The authors have declared no competing interest.

