## Supplementary Materials for "Localized modulation of DNA supercoiling, triggered by the *Shigella* anti-silencer VirB, is sufficient to relieve H-NS-mediated silencing"

**Table of Contents**

Supplemental Materials and Methods 2

Table S1. Bacterial strains and plasmids used in this study 3

Table S2. Primers and DNA Duplexes used in this study 5

Table S3. Complete statistics for *topA* mutant assays 8

Table S4. Complete statistics for assay exploiting twin domain model of transcription 9

Supplementary Figures with Legends 10

References 22

### Supplemental Materials and Methods

#### Plasmid construction

For the construction of the mutated pBAD-*virB* producing VirB protein with K152E amino acid substitution (pADK15), a two-step, mutagenic megaprimer PCR method was used. First, W651 & 652 were used to generate the mutagenic megaprimer (214 bp) using pATM324 (pBAD-*virB*) as a template. The mutagenic megaprimer generated from the first PCR reaction was then used with W653 to amplify a 704 bp amplicon from pATM324. The resulting amplicon was digested with *Bgl*II and *Hind*III restriction enzymes and ligated into pATM324 digested with the same restriction enzymes.

For the construction of the L-arabinose inducible pT7-RNAP (pMK26), *gene1* encoding the T7 RNA polymerase was digested with *Sac*I and *Xba*I pTARA ((1); Addgene # 31491) and ligated into pBAD18 (2) digested with the same restriction enzymes. L-arabinose-dependent production of T7 RNAP was verified using SDS-PAGE (data not shown). To make a promoterless control (pMK29) G-block #14, carrying a multi-cloning site from pBS, was digested with *Pst*I and *Xba*I and ligated into pAFW04a digested with the same restriction enzymes.

#### Strain Verification

Before use, the *topA* alleles in MG1655 (NCBI reference NC_000913.3), DPB923, and DPB924 were compared by Sanger sequencing (each strain received from the *E. coli* stock center). Analyses revealed that the *topA10* allele in DPB923 contained several missense mutations before a premature stop. As such, the protein is 68 amino acids shorter (R790G, E791D, T792A, R793A, A794N, P795L, V797_K865delinsIX) than the wild type allele found in MG1655 and DPB924. To the best of our knowledge, these DNA sequence details of the *topA10* alleles are the first to be reported and corroborate Western analyses of these strains by Sternglanz *et al*., 1981 (3), where they concluded *topA10* (DPB923) encoded a truncated version of the DNA Topoisomerase I protein.

#### Promoter activity assays with novobiocin

The *Shigella flexneri virB* mutant AWY3 carrying pAFW04 was grown in triplicate overnight at 30 °C with shaking in LB containing a final concentration of chloramphenicol at 20 μg $ml^{-1}$. Cells were diluted 1:100 in fresh LB with chloramphenicol and grown for 3 hours at 37 °C. Cultures were normalized to an OD600 of 0.003 (4) and novobiocin was immediately added at 0, 15, or 25 μg $ml^{-1}$ final concentration. Cells were grown for an additional 3 hours at 37 °C with shaking before harvesting. The β-galactosidase activities were determined as described previously (5).

**Table S1. Bacterial strains and plasmids used in this study**

| Label | Description | Reference |
|---|---|---|
| Strains | | |
| *S. flexneri* | | |
| 2457T | Wild-type *S. flexneri* serotype 2a | (6) |
| AWY3 | 2457T *virB*::Tn5; $Kn^r$ | (5) |
| *E. coli* | | |
| DH10B | $F^-$ *endA1 deoR*$^+$ *recA1 galE15 galK16 nupG rpsL* Δ(*lac*)*X74 φ80lacZ*Δ*M15 araD139* Δ(*ara, leu*)*7697 mcrA* Δ(*mrr-hsdRMS-mcrBC*) *Str*$^R$ $λ^-$ | (7) |
| MC4100 | F-, *[araD139]B/r,* Δ*(argF-lac)169,* λ-, *e14-, flhD5301,* Δ*(fruK-yeiR)725(fruA25), relA1, rpsL150(strR), rbsR22,* Δ*(fimB-fimE)632(::IS1), deoC1* | (8) |
| MC4100 *hns*::$Kn^r$ | *hns* dominant negative mutant expressing 1 - 37 amino acids | (9) |
| DH10B *hns*::$Kn^r$ | DH10B with *hns*::$Kn^r$ allele transduced from MC4100 *hns*::$Kn^r$ | This Work |
| MG1655 | Wildtype *E. coli* F-, λ-, *rph-1* | (10) CGSC# 6300 |
| DPB923 | *topA10* mutant F-, λ-, *topA10, rph-1* | (11) CGSC# 7894 |
| DBP924 | *topA10*$^+$ F-, λ-, *rph-1* | (11) CGSC# 7893 |
| Plasmids | | |
| pHJW20 | P*icsP-lacZ* reporter plasmid derived from pACYC184 | (5) |
| pMIC21 | pHJW20 lacking all promoter sequences | (12) |
| pCTH03 | pHJW20 truncated from the 3' end to -32 relative to the TSS; remove P1 and P2 | (13) |
| pMIC18 | pHJW20 with VirB boxes 1 and 2 mutated by transition mutations | (12) |

| | | |
|---|---|---|
| pMAP27 | pHJW20 with *icsP* sequences downstream of -426 relative to TSS and entire lacZ gene removed; promoter-proximal H-NS binding region and *lacZ* gene excised. | This Work |
| pMAP29 | pMAP27 with *icsP* sequences downstream of -964 relative to TSS through +27 of entire lacZ gene removed; promoter-distal and proximal H-NS binding regions and *lacZ* gene excised. | This Work |
| pMK63 (pDivergent) | pHJW20 with P*phi10*-*gfp2*-5xT inserted at -1226 upstream and in the opposite direction of P*icsP* | This Work |
| pMK64 (pParallel) | pHJW20 with P*phi10*-*gfp2*-5xT inserted -1226 upstream and in the same direction of P*icsP* | This Work |
| pLL39 | Source of 5x *rrnB* T1 terminator | (14) Addgene #15458 |
| pAFW04 | Identical to pHJW20, but the lambda oop terminator is cloned immediately upstream of the *icsP* promoter fragment to prevent possible transcriptional read-through into the promoter region. | (15) |
| pMK29 | pAFW04 with a MCS from pBlueScript II KS+ replacing the entire 1259 bp *icsP* promoter to make a promoterless control. | This Work |
| pBAD18 | Arabinose-inducible pBAD expression vector, pBR ori; Ampr$^{R}$ | (2) |
| pMK26 | pMK26 is pBAD18-CmR expressing T7 RNAP sourced from pTARA (1) using SacI and XbaI. | This Work |
| pTARA | T7 RNAP expression plasmid in a pBAD33 vector. | (1) |
| pATM324 | *virB* expression plasmid in a pBAD18 vector | (16) |
| pADK15 | pATM324 derivative that produces VirB K152E | This Work |

**Table S2. Primers and DNA Duplexes used in this study**

| Primer | Sequence 5’ to 3’ | Description and Use |
|---|---|---|
| W43 | CTCTACTGTTTCTCCATACCC | Used for sequencing pATM324 derivative; binds 60 bp upstream of ATG |
| W93 | TGGGTTGAAGGCTCTCAAGGGC | Used to sequence pMK29 |
| W119 | GCCAGGGTTTTCCCAGTCACGA | Used to sequence Gblock 13 carried in pBlueScript SK II+ |
| W391 | /5Phos/GATTGAATACTTCCGGGGATTTCAGTATGAAATGAAGTATATTTAATATACTTT | Top strand to be annealed with W392; Target 6 wild-type VirB boxes; 54 bp |
| W392 | AAAGTATATTAAATATACTTCATTTCATACTGAAATCCCCGGAAGTATTCAATC | Bottom strand to be annealed with W392; Target 6 wild-type VirB boxes; 54 bp |
| W393 | GATTGAATACTTCCGGGGcgggtcgTgctgggcGAAGTATATTTAATATACTTT | Bottom strand to be annealed with W392; Target 6 mutated VirB boxes (lowercase); 54 bp |
| W394 | /5Phos/GATTGAATACTTCCGGGGgcccagcTcgacccgGAAGTATATTTAATATACTTT | Top strand to be annealed with W392; Target 6 mutated VirB boxes (lowercase); 54 bp |
| W428 | ATTGCACAACTGAATTTAAGGC | Used with W429 to amplify *hns* locus; binds 50 bp upstream of ATG |
| W429 | TTAAATTGTCTTAAACCG | Used with W428 to amplify *hns* locus; binds 56 bp downstream of TAA |
| W651 | CGAGACAGATTCTCTTTTTTGGCgATaTCcTcATAGGACATCCC | Mutagenic primer codon 152 of *virB* ORF; TTT–>**cTc** to give K152E; *Eco*RV created by silent mutations (lowercase); Used with W652 to generate mutagenic megaprimer |
| W652 | GCACTCGTAGAAGAGCATCTGCA | Used with W651 to generate mutagenic megaprimer; binds 272 bp downstream of *virB* ATG |

| | | |
|---|---|---|
| W653 | GGCTGAAAATCTTCTCTCATCCGCC | Used mutagenic megaprimer to generate insert for pADK15; binds 46 bp downstream of *virB* TAA |
| W765 | GCAGGAGTCGCATAAGGG | Used to sequence Gblock 13 carried in pBlueScript SK II+ |
| W766 | GACAGTCATAAGTGCGGCG | Used to sequence Gblock 13 carried in pBlueScript SK II+ |
| W767 | GCTCACTCATTAGGCACCCC | Used to sequence Gblock 13 carried in pBlueScript SK II+ |
| W800 | ACTGATGCTAGCAGGAGGAATTCACCATGAGTAAAGGAGAAGAACTTTT<br>CACTGG | Used to amplify *gfp* replacing *tetA* in Gblock13 |
| W801 | ATACGTGACGTCTCATTATTTGTATAGTTCATCCATGCC | Used to amplify *gfp* replacing *tetA* in Gblock13 |
| W864 | GGTTACGAGATCGAAGAGGGCGAATTCCGC | Verify *topA 10* locus in MG1655, DPB923 and DPB924 |
| P*φ10-gfp-2-*5xT<br><br>insert | ATAGGCACTGCAGATAGATTACAGCTGGCTTCCGGCTCGTATGTTGTGT<br>GGAATTGTGAGCGGATAACAATTTCACACAGGAAACAGCTATGACCATG<br>ATTACGAATT**TAATACGACTCACTATAGGGA**TTCTCCATACCCGTTTTT<br>TTGGGCTAGCAGGAGGAATTCACC**ATGAGTAAAGGAGAAGAACTTTTCA**<br>**CTGGAGTTGTCCCAATTCTTGTTGAATTAGATGGTGATGTTAATGGGCA**<br>**CAAATTTTCTGTCAGTGGAGAGGGTGAAGGTGATGCAACATACGGAAAA**<br>**CTTACCCTTAAATTTATTTGCACTACTGGAAAACTACCTGTTCCATGGC**<br>**CAACACTTGTCACTACTTTCGCGTATGGTCTTCAATGCTTTGCGAGATA**<br>**CCCAGATCATATGAAACAGCATGACTTTTTCAAGAGTGCCATGCCCGAA**<br>**GGTTATGTACAGGAAAGAACTATATTTTTCAAAGATGACGGGAACTACA**<br>**AGACACGTGCTGAAGTCAAGTTTGAAGGTGATACCCTTGTTAATAGAAT**<br>**CGAGTTAAAAGGTATTGATTTTAAAGAAGATGGAAACATTCTTGGACAC**<br>**AAATTGGAATACAACTATAACTCACACAATGTATACATCATGGCAGACA**<br>**AACAAAAGAATGGAATCAAAGTTAACTTCAAAATTAGACACAACATTGA**<br>**AGATGGAAGCGTTCAACTAGCAGACCATTATCAACAAAATACTCCAATT**<br>**GGCGATGGCCCTGTCCTTTTACCAGACAACCATTACCTGTCCACACAAT**<br>**CTGCCCTTTCGAAAGATCCCAACGAAAAGAGAGACCACATGGTCCTTCT**<br>**TGAGTTTGTAACAGCTGCTGGGATTACACATGGCATGGATGAACTATAC**<br>**AAATAA**TGAGACGTCTAAGAAACCATTATTATCATGGGGTATCGATAAG<br>CTTGATATCGAATTCACTAGTGATTGGGCCCATTACAGCGGATCAATTC | Insert carrying the P*φ10-gfp-2-5xT* to be used make pMK63 and pMK64<br><br>Notable restriction sites are underlined and P*φ10, gfp-2,* and the 5x *rrnB* T1 terminators are all **bolded** |

| | | |
|---|---|---|
| | CCAATTCCCCAGGCATCAA**ATAAAACGAAAGGCTCAGTCGAAAGACTGG**<br>**GCCTTTCGTTTTATCTGTTGTTTGTCGGTGAACGCTCTCCTGAGTAGGA**<br>**CAAATCCGCCGGGAGCGGATTTGAACGTTGCGAAGCAACGGCCCGGAGG**<br>**GTGGCGGGCAGGACGCCCGCCATAAACTGCCAGGAATTAATTCCCCAGG**<br>**CATCAAATAAAACGAAAGGCTCAGTCGAAAGACTGGGCCTTTCGTTTTA**<br>**TCTGTTGTTTGTCGGTGAACGCTCTCCTGAGTAGGACAAATCCGCCGGG**<br>**AGCGGATTTGAACGTTGCGAAGCAACGGCCCGGAGGGTGGCGGGCAGGA**<br>**CGCCCGCCATAAACTGCCAGGAATTAATTCCCCAGGCATCAAATAAAAC**<br>**GAAAGGCTCAGTCGAAAGACTGGGCCTTTCGTTTTATCTGTTGTTTGTC**<br>**GGTGAACGCTCTCCTGAGTAGGACAAATCCGCCGGGAGCGGATTTGAAC**<br>**GTTGCGAAGCAACGGCCCGGAGGGTGGCGGGCAGGACGCCCGCCATAAA**<br>**CTGCCAGGAATTAATTCCCCAGGCATCAAATAAAACGAAAGGCTCAGTC**<br>**GAAAGACTGGGCCTTTCGTTTTATCTGTTGTTTGTCGGTGAACGCTCTC**<br>**CTGAGTAGGACAAATCCGCCGGGAGCGGATTTGAACGTTGCGAAGCAAC**<br>**GGCCCGGAGGGTGGCGGGCAGGACGCCCGCCATAAACTGCCAGGAATTA**<br>**ATTCCCCAGGCATCAAATAAAACGAAAGGCTCAGTCGAAAGACTGGGCC**<br>**TTTCGTTTTATCTGTTGTTTGTCGGTGAACGCTCTCCTGAGTAGGACAA**<br>**ATCCGCCGGGAGCGGATTTGAACGTTGCGAAGCAACGGCCCGGAGGGTG**<br>**GCGGGCAGGACGCCCGCC**ATAAACTGCCAGGAATTGGGGATCGGAATTC<br>CCATATGGACTCAGCTGCATGGGATAAAGCTTATGGAAC | |
| Gblock 14 | ATATACTGCAGGCGAATTGGAGCTCCACCGCGGTGGCGGCCGCACTAGT<br>GGATCCCCCGGGGAATTCGATATCAAGCTTATCGATACCGTCGACCTCG<br>AGGGGGGGCCCGGTACCCAGCTTTTGTTCCCTTTAGTGATCTAGAATAT<br>A | DNA carrying a multicloning site from pBlueScript KS II+, used to make pMK29 |

**Table S3. Complete statistics for *topA* mutant assays**

| ***topA* mutant assay** | strain | **MG1655** | | ***topA10*** | | ***top+*** | |
|---|---|---|---|---|---|---|---|
| strain | plasmids | **pEmpty** | **pBAD-*virB*** | **pEmpty** | **pBAD-*virB*** | **pEmpty** | **pBAD-*virB*** |
| **MG1655** | **pEmpty** | | 0.001* | 1 | | 1 | |
| | **pBAD-*virB*** | | | | 0.001* | | 0.045* |
| ***topA10*** | **pEmpty** | | | | 0.001* | 1 | |
| | **pBAD-*virB*** | | | | | | 0.001* |
| ***top+*** | **pEmpty** | | | | | | 0.001* |
| | **pBAD-*virB*** | | | | | | |

Significance was determined using a two-way ANOVA with post-hoc Bonferroni. Asterisks indicate p<0.05. Grey boxes represent data that was not compared.

**Table S4. Complete statistics for assays exploiting twin domain model of transcription**

| **GFP fluorescence** | | strain | **MC4100** | | | | | | ***hns::kn^R*** | | | | | |
|---|---|---|---|---|---|---|---|---|---|---|---|---|---|---|
| | | pBAD derivative | **pBAD18** | | | **pT7RNAP** | | | **pBAD18** | | | **pT7RNAP** | | |
| strain | pBAD derivative | reporter | **P*icsP-lacZ*** | **pDivergent** | **pParallel** | **P*icsP-lacZ*** | **pDivergent** | **pParallel** | **P*icsP-lacZ*** | **pDivergent** | **pParallel** | **P*icsP-lacZ*** | **pDivergent** | **pParallel** |
| **MC4100** | **pBAD18** | **P*icsP-lacZ*** | | 1 | 0.665 | 0.901 | | | 0.435 | | | | | |
| | | **pDivergent** | | | 0.376 | | 0.001* | | | 0.891 | | | | |
| | | **pParallel** | | | | | | 0.001* | | | 0.123 | | | |
| | **pT7RNAP** | **P*icsP-lacZ*** | | | | | 0.001* | 0.001* | | | | 0.525 | | |
| | | **pDivergent** | | | | | | 0.001* | | | | | 0.001* | |
| | | **pParallel** | | | | | | | | | | | | 0.638 |
| ***hns::kn^R*** | **pBAD18** | **P*icsP-lacZ*** | | | | | | | | 1 | 1 | 0.981 | | |
| | | **pDivergent** | | | | | | | | | 1 | | 0.001* | |
| | | **pParallel** | | | | | | | | | | | | 0.001* |
| | **pT7RNAP** | **P*icsP-lacZ*** | | | | | | | | | | | 0.001* | 0.001* |
| | | **pDivergent** | | | | | | | | | | | | 0.001* |
| | | **pParallel** | | | | | | | | | | | | |

| ***lacZ* expression** | | strain | **MC4100** | | | | | | ***hns::kn^R*** | | | | | |
|---|---|---|---|---|---|---|---|---|---|---|---|---|---|---|
| | | pBAD derivative | **pBAD18** | | | **pT7RNAP** | | | **pBAD18** | | | **pT7RNAP** | | |
| strain | pBAD derivative | reporter | **P*icsP-lacZ*** | **pDivergent** | **pParallel** | **P*icsP-lacZ*** | **pDivergent** | **pParallel** | **P*icsP-lacZ*** | **pDivergent** | **pParallel** | **P*icsP-lacZ*** | **pDivergent** | **pParallel** |
| **MC4100** | **pBAD18** | **P*icsP-lacZ*** | | 1 | 1 | 0.691 | | | 0.001* | | | | | |
| | | **pDivergent** | | | 1 | | 0.284 | | | 0.001* | | | | |
| | | **pParallel** | | | | | | 0.001* | | | 0.001* | | | |
| | **pT7RNAP** | **P*icsP-lacZ*** | | | | | 0.76 | 0.001* | | | | 0.001* | | |
| | | **pDivergent** | | | | | | 0.76 | | | | | 0.006* | |
| | | **pParallel** | | | | | | | | | | | | 0.001* |
| ***hns::kn^R*** | **pBAD18** | **P*icsP-lacZ*** | | | | | | | | 0.756 | 1 | 0.968 | | |
| | | **pDivergent** | | | | | | | | | 0.281 | | 0.001* | |
| | | **pParallel** | | | | | | | | | | | | 0.001* |
| | **pT7RNAP** | **P*icsP-lacZ*** | | | | | | | | | | | 0.001* | 0.001* |
| | | **pDivergent** | | | | | | | | | | | | 0.001* |
| | | **pParallel** | | | | | | | | | | | | |

Significance was determined using a three-way ANOVA with post-hoc Bonferroni. Asterisks indicate $p<0.05$. Grey boxes represent data that was not compared.

**Supplemental Figures with Legends**

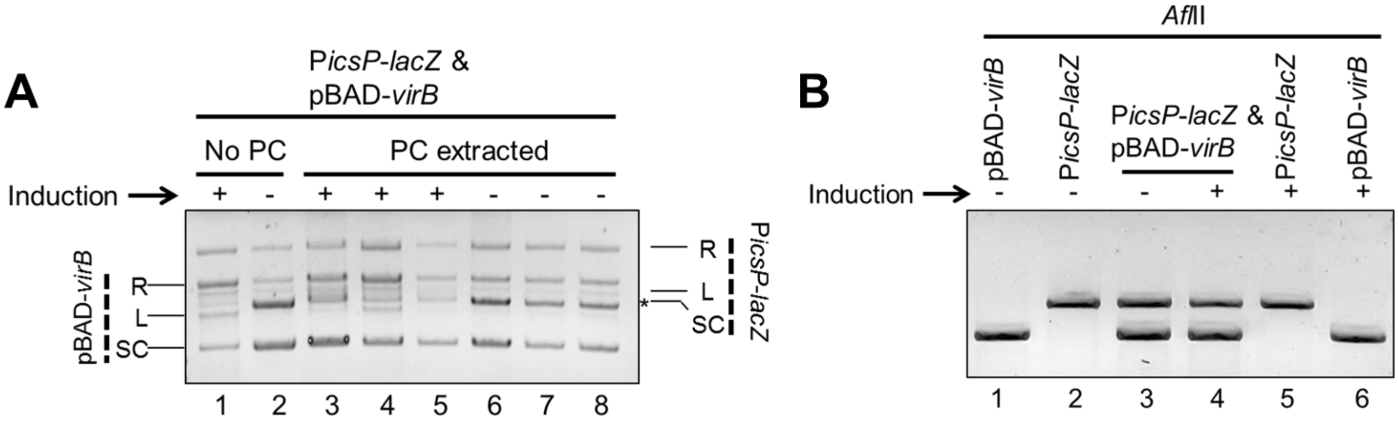


**Figure S1. Electrophoretic mobility of P*icsP-lacZ*, pBAD-*virB* or both isolated from *E. coli* DH10B with or without induction.** A) Electrophoresis after additional phenol-chloroform extraction to assess if altered mobility was due to residual protein binding. B) Electrophoresis of samples restricted with *Afl*II to assess if the altered mobility of the P*icsP-lacZ* reporter was triggered by a VirB-dependent change in its size.

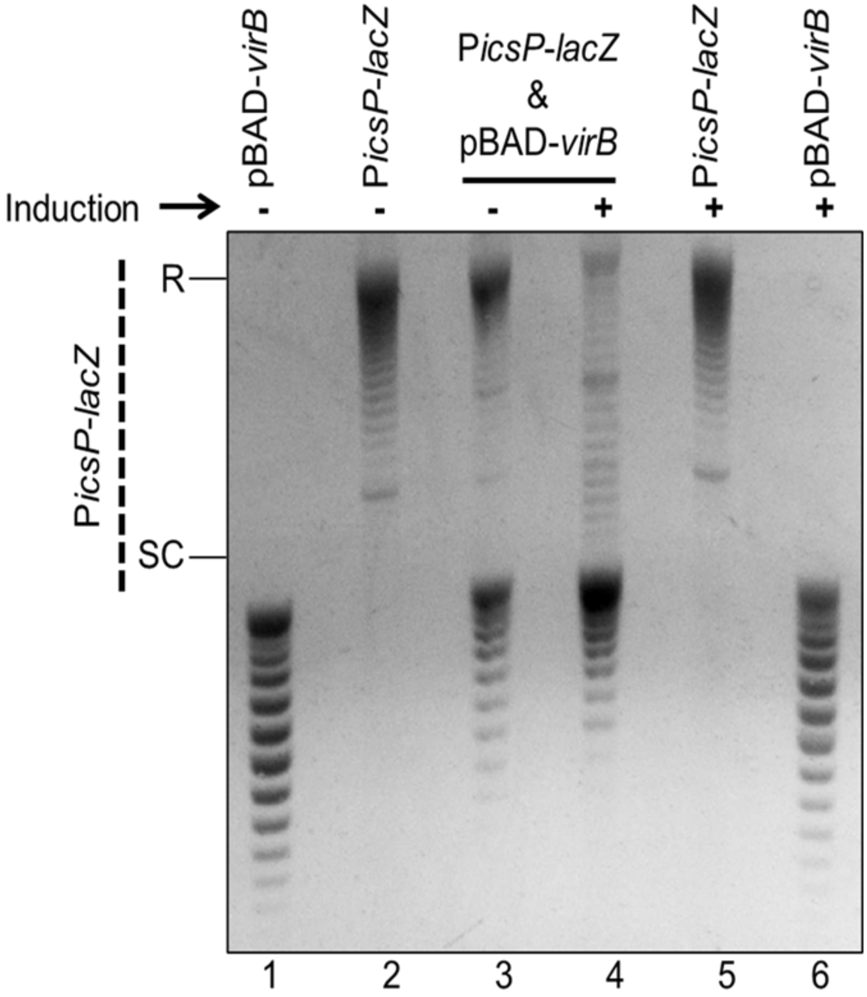


**Figure S2. Topoisomer analysis of identically isolated samples from Figure 1C analysed on an agarose gel containing higher concentration of chloroquine (10 μg ml$^{-1}$).** Under these conditions negatively supercoiled DNA became more relaxed and migrated less during electrophoresis (Figure S2A, lanes 1-3, 5 & 6 and pBAD-*virB* in lane 4) whereas the *PicsP-lacZ* reporter in the DNA samples containing two plasmids isolated from cells expressing *virB* (Figure S2A, lane 4)*,* migrated faster (Figure S1C; compare lane 3 with 4). These findings are consistent with the P*icsP-lacZ* reporter isolated from cells with VirB being less negatively supercoiled. R = relaxed, SC = supercoiled. Quantification of key lanes (3 & 4) is provided in Figure S6 E & F.

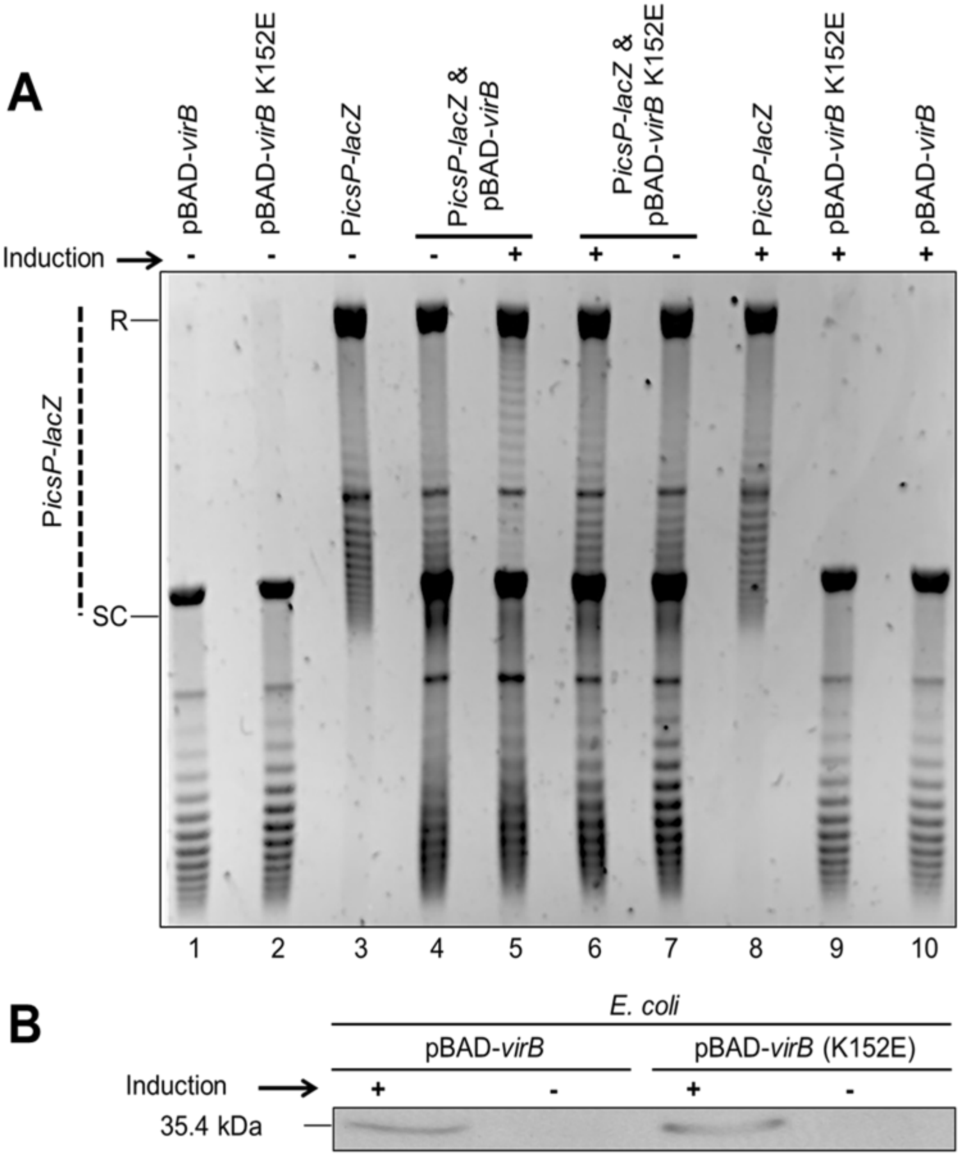


**Figure S3. Analysis of the role of DNA binding in VirB-dependent changes in DNA supercoiling of P*icsP-lacZ*.**
A) Topoisomer analysis of the wild-type P*icsP-lacZ* reporter isolated in the presence of pBAD expressing either wild-type VirB or VirB K152E deficient in DNA binding (17,18) after electrophoresis in the presence of chloroquine (2.5 μg $ml^{-1}$). Quantification of key lanes (4-7) is provided in Figure S6, M-R. B) Western blot of wild-type VirB and VirB K152E (17,18) prepared from *E. coli* in the presence or absence of induction. Taken together, data show the DNA binding mutant causes a less pronounced the change in topoisomer distribution, even though VirB and VirB K152E are produced at equivalent levels under the conditions used.

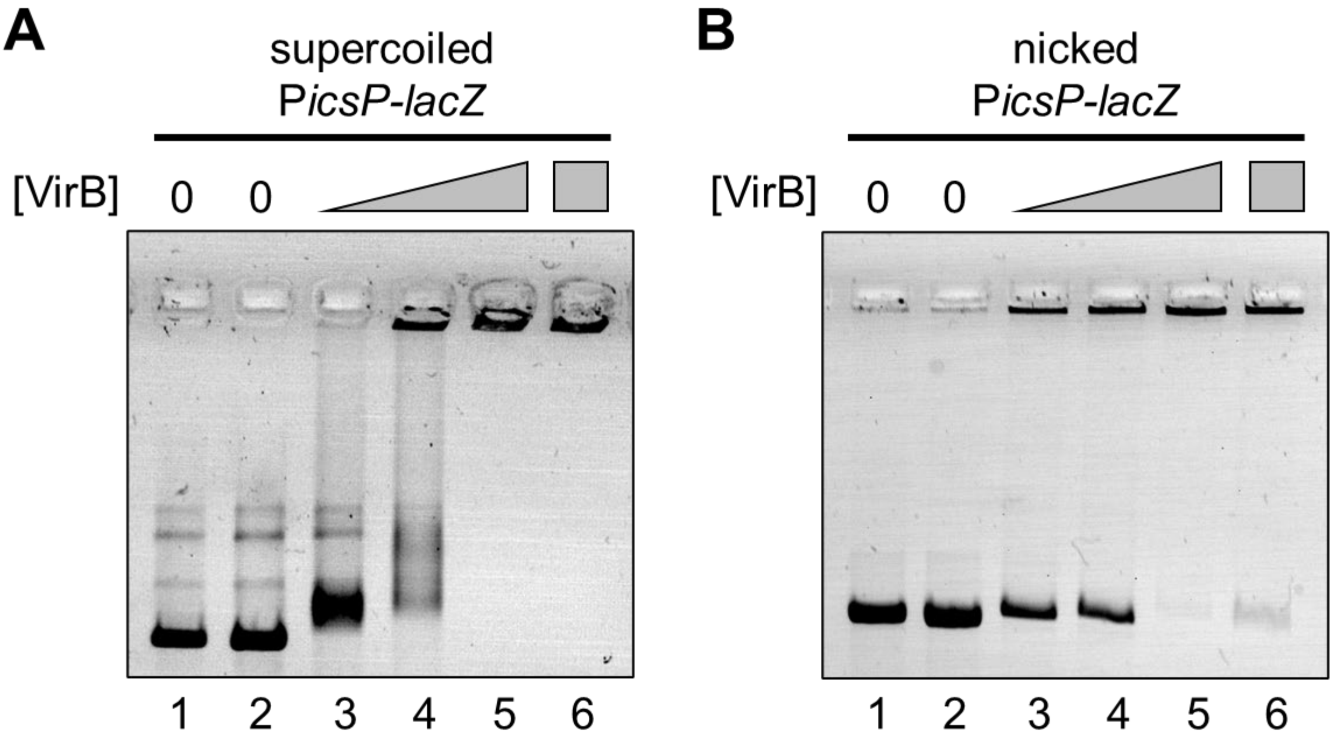


**Figure S4. Electrophoretic mobility shift assays that accompany the *in vitro* assays shown in Figure 6. A)** Electrophoretic mobility of negatively supercoiled P*icsP-lacZ* plasmid (3.3 nM) incubated with increasing concentrations of VirB (lanes 1 and 2= 0 μM, lane 3 = 0.12 μM, lane 4 = 0.35 μM, lanes 5 and 6 = 0.59 μM). **B)** Electrophoretic mobility of nicked P*icsP-lacZ* plasmid (3.0 nM) with increasing concentrations of VirB (lanes 1 and 2 = 0 μM, lane 3 = 0.12 μM, lane 4 = 0.24 μM, lanes 5 and 6 = 0.35 μM).

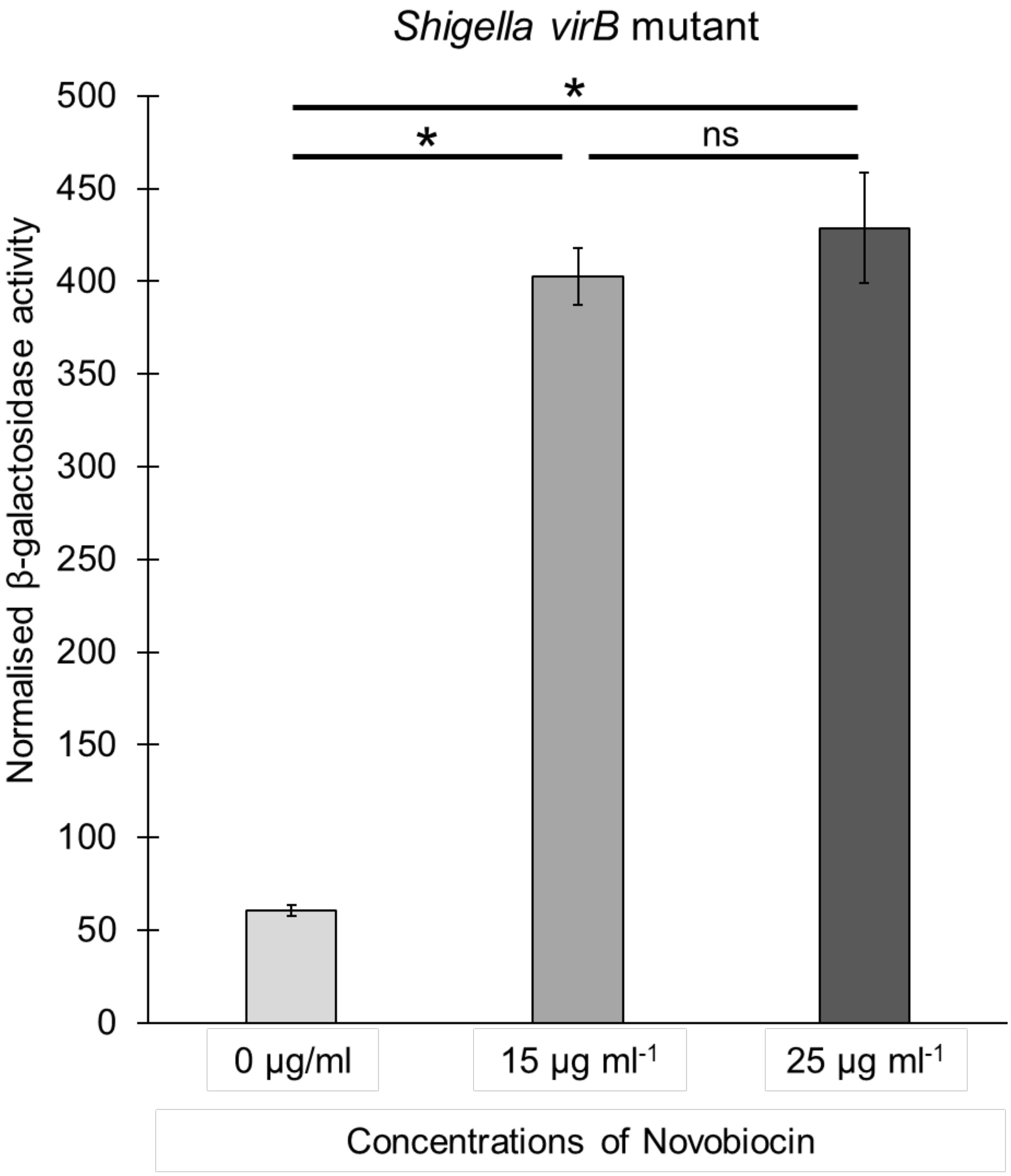


**Figure S5. Effect of novobiocin on *icsP* promoter activity as judged by β-galactosidase assays.** Normalised promoter activity of P*icsP*-*lacZ* measured in *Shigella virB* mutant exposed to 0, 15, or 25 µg $ml^{-1}$ novobiocin. Data are consistent with the accrual of positive supercoils relieving H-NS-mediated silencing of P*icsP*, leading to enhanced P*icsP*-*lacZ* activity. Assays were done with three biological replicates and repeated three times. Representative data are shown. Significance was calculated using a one-way ANOVA, with post-hoc Bonferroni, *, $p<0.05$ and ns, not significant.

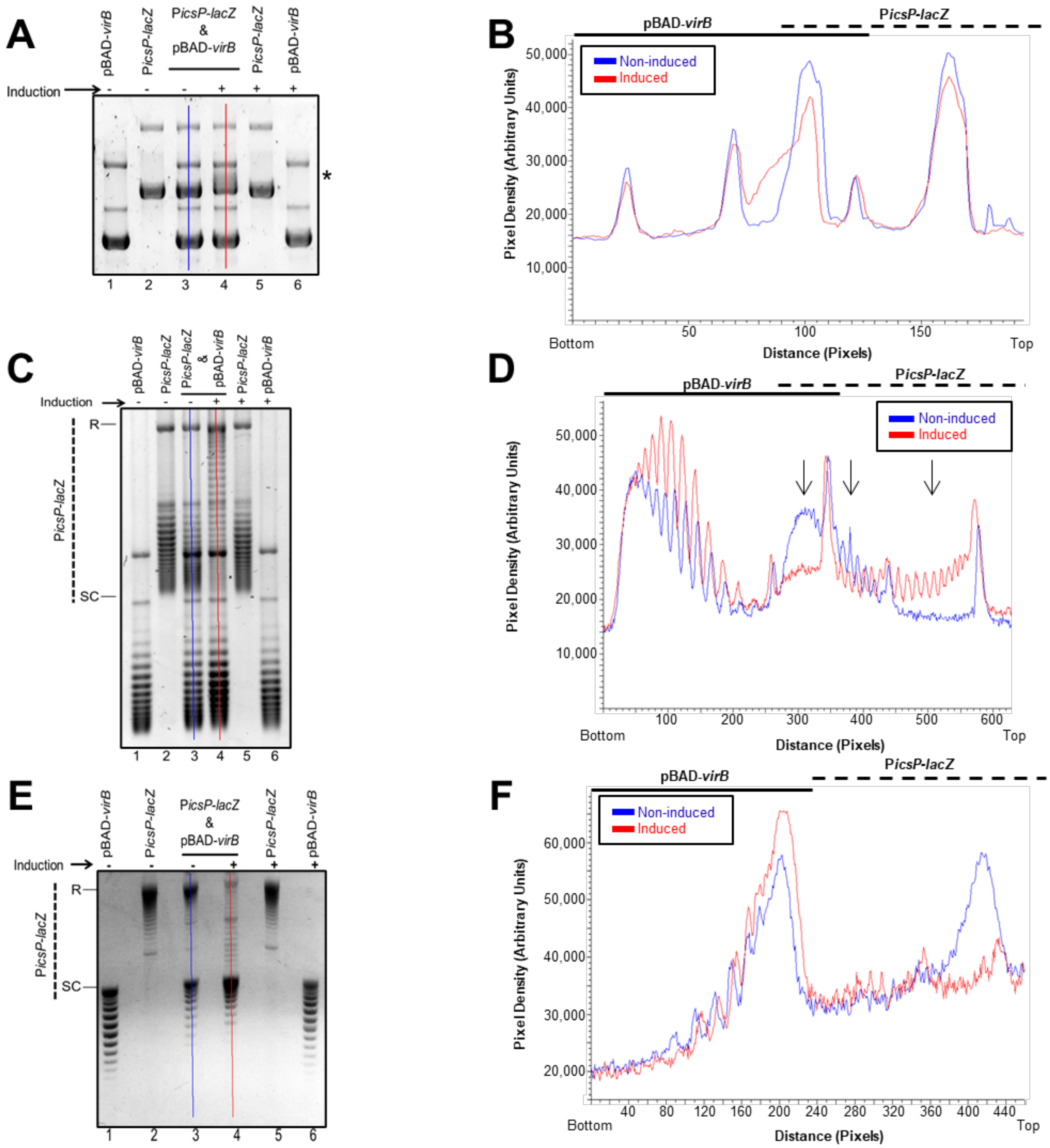
A
pBAD-virB
PicsP-lacZ
PicsP-lacZ & pBAD-virB
Induction
*
B
pBAD-virB
PicsP-lacZ
Non-induced
Induced
Pixel Density (Arbitrary Units)
Distance (Pixels)
Bottom
Top
C
R
SC
D
E
F

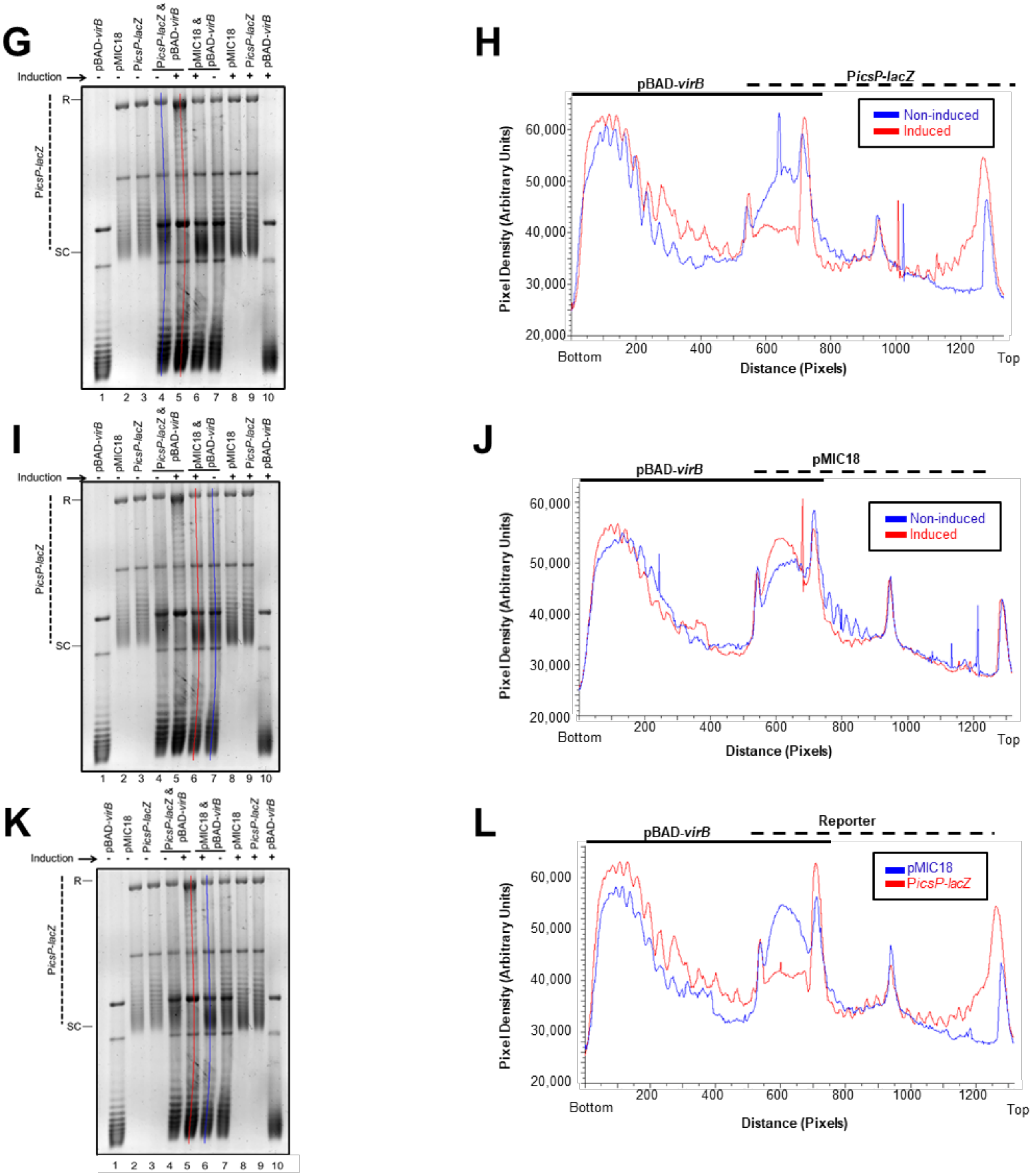
G
H
I
J
K
L
pBAD-virB
pMIC18
PicsP-lacZ
PicsP-lacZ & pBAD-virB
pMIC18 & pBAD-virB
Induction
R
SC
1 2 3 4 5 6 7 8 9 10
Non-induced
Induced
Pixel Density (Arbitrary Units)
Distance (Pixels)
Bottom
Top
Reporter
60,000
50,000
40,000
30,000
20,000
200
400
600
800
1000
1200

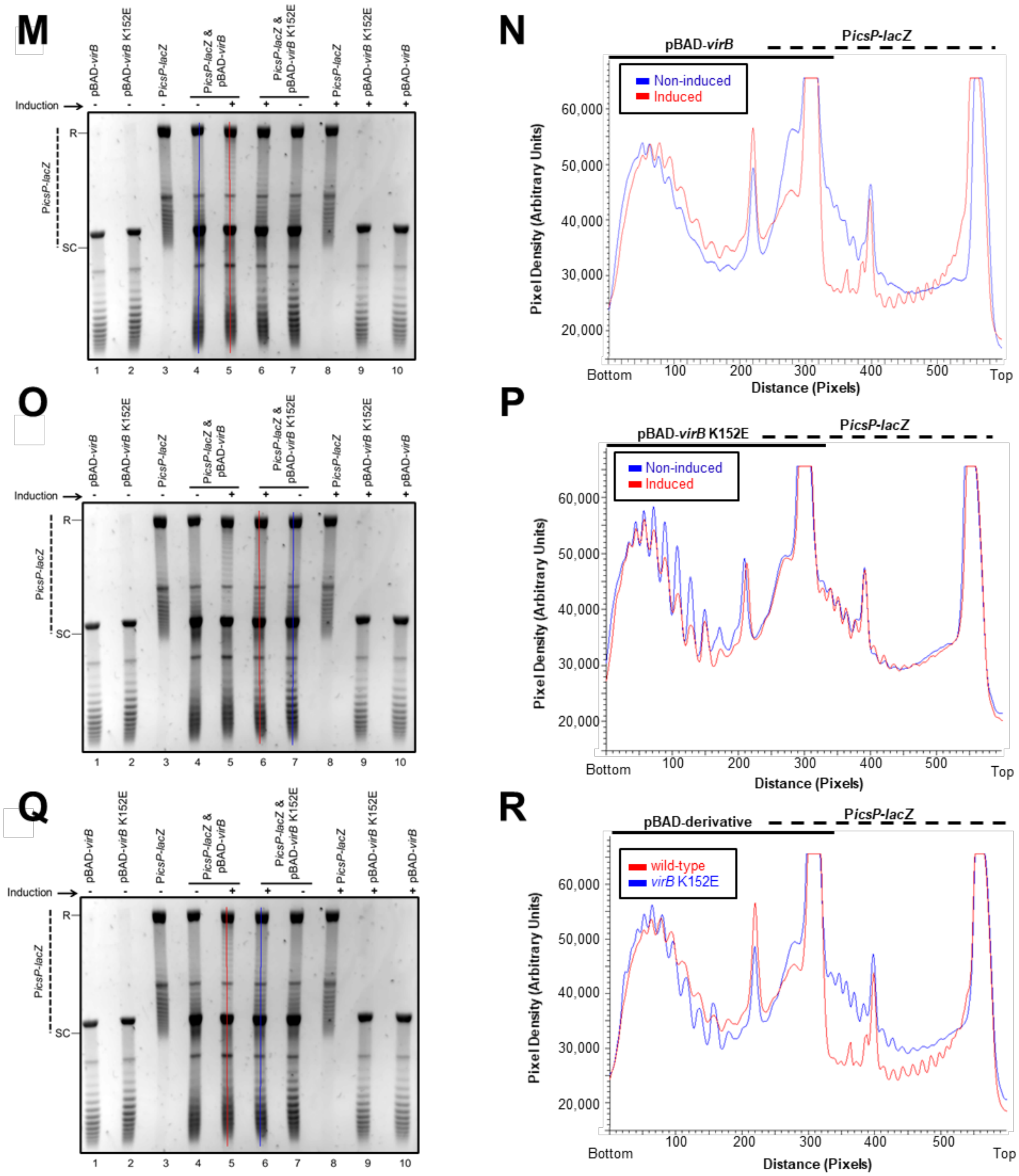
M
pBAD-virB
pBAD-virB K152E
PicsP-lacZ
PicsP-lacZ & pBAD-virB
PicsP-lacZ & pBAD-virB K152E
PicsP-lacZ
pBAD-virB K152E
pBAD-virB
Induction
R
SC
PicsP-lacZ
1 2 3 4 5 6 7 8 9 10
N
pBAD-virB
PicsP-lacZ
Non-induced
Induced
Pixel Density (Arbitrary Units)
20,000
30,000
40,000
50,000
60,000
Bottom
100
200
300
400
500
Top
Distance (Pixels)
O
P
pBAD-virB K152E
Q
R
pBAD-derivative
wild-type
virB K152E

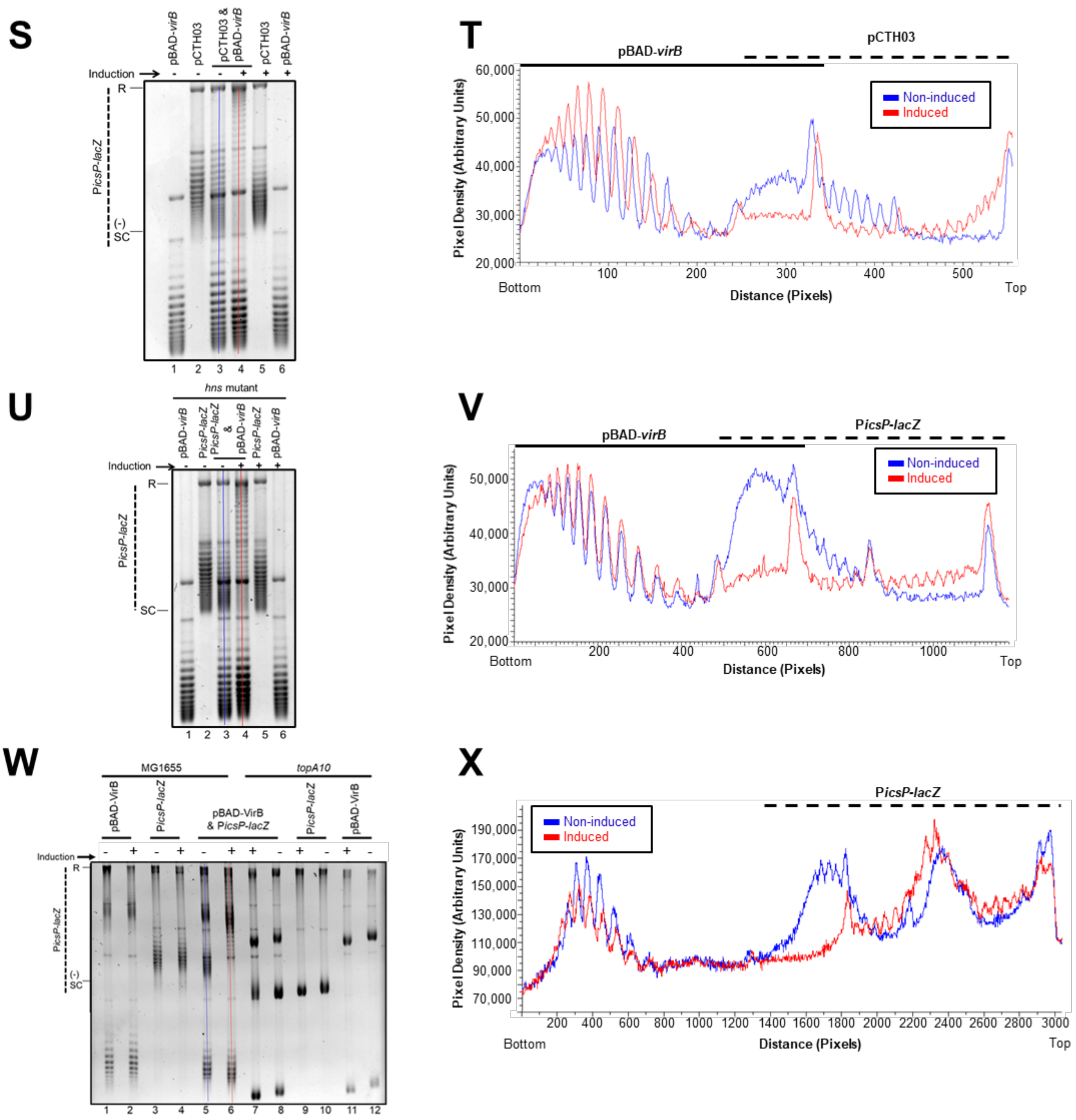

S
Induction
pBAD-virB
pCTH03
pCTH03 & pBAD-virB
pCTH03
pBAD-virB
R
PicsP-lacZ
(-) SC
1 2 3 4 5 6
T
pBAD-virB
pCTH03
Non-induced
Induced
Pixel Density (Arbitrary Units)
Bottom
Distance (Pixels)
Top
U
hns mutant
Induction
pBAD-virB
PicsP-lacZ
PicsP-lacZ & pBAD-virB
PicsP-lacZ
pBAD-virB
R
PicsP-lacZ
SC
1 2 3 4 5 6
V
pBAD-virB
PicsP-lacZ
Non-induced
Induced
Pixel Density (Arbitrary Units)
Bottom
Distance (Pixels)
Top
W
MG1655
topA10
pBAD-VirB
PicsP-lacZ
pBAD-VirB & PicsP-lacZ
PicsP-lacZ
pBAD-VirB
Induction
R
PicsP-lacZ
(-) SC
1 2 3 4 5 6 7 8 9 10 11 12
X
PicsP-lacZ
Non-induced
Induced
Pixel Density (Arbitrary Units)
Bottom
Distance (Pixels)
Top

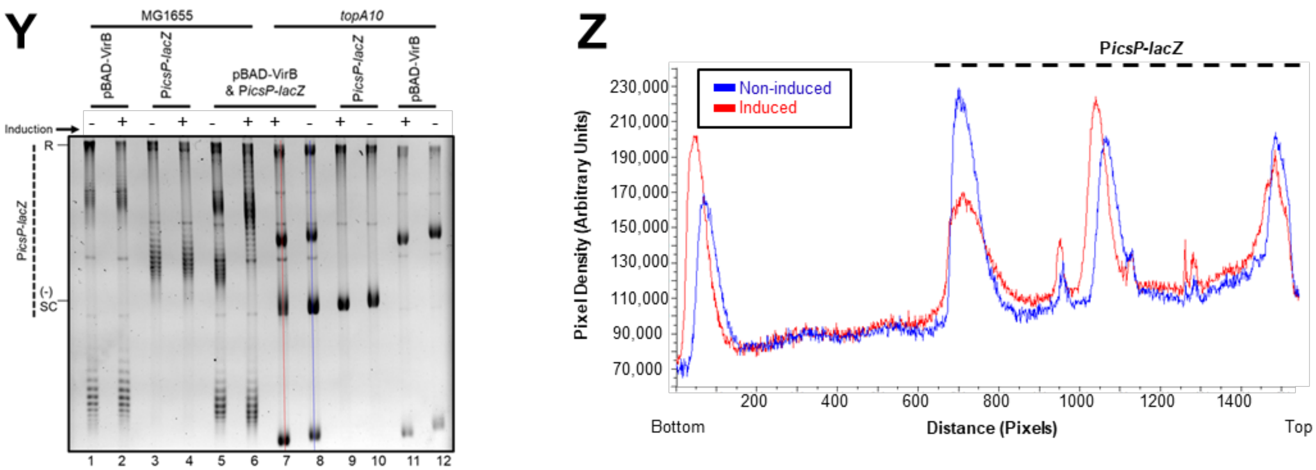


**Figure S6. Quantification of DNA isolated from cells after 1D gel electrophoresis.** Lane trace analyses are shown in panels on the left. Lane traces were routinely drawn from the bottom of the gel to the top. The profile of lane traces is plotted in corresponding panels on the right; solid lines denote signals corresponding to pBAD-*virB* or derivatives, and dashed lines denote the signal corresponding to P*icsP-lacZ* or derivatives. In each case, lane trace analyses of two lanes on the respective gel is shown, facilitating comparison of respective lanes without signal crowding. Panels A-B relate to Figure 1B, C-D relate to Figure 1C, E-F relate to Figure S2, G-L relate to Figure 2, M-R relate to Figure S3A, S-T relate to Figure 3A, U-V relate to Figure 3B and W-Z relate to Figure 5A. For ease of interpretation, major changes between topoisomer distributions in P*icsP-lacZ* in the presence/absence of *virB* are highlighted with downward facing arrows in panel D, the initial panel where these changes were observed.

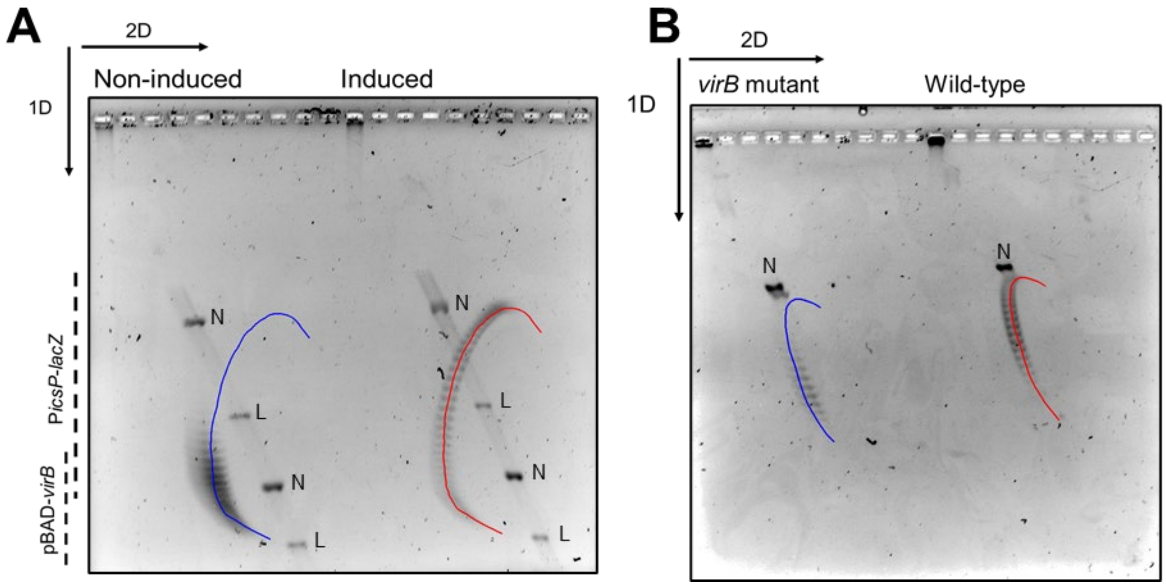


**Figure S7. Arcs used for 2D gel analysis.** A) Arcs drawn for 2D gel analysis shown in Figure 4C, red = induced and blue = non-induced. B) Arcs drawn for 2D gel analysis shown in Figure 4E. blue = *virB* mutant and red = wild-type *virB*.

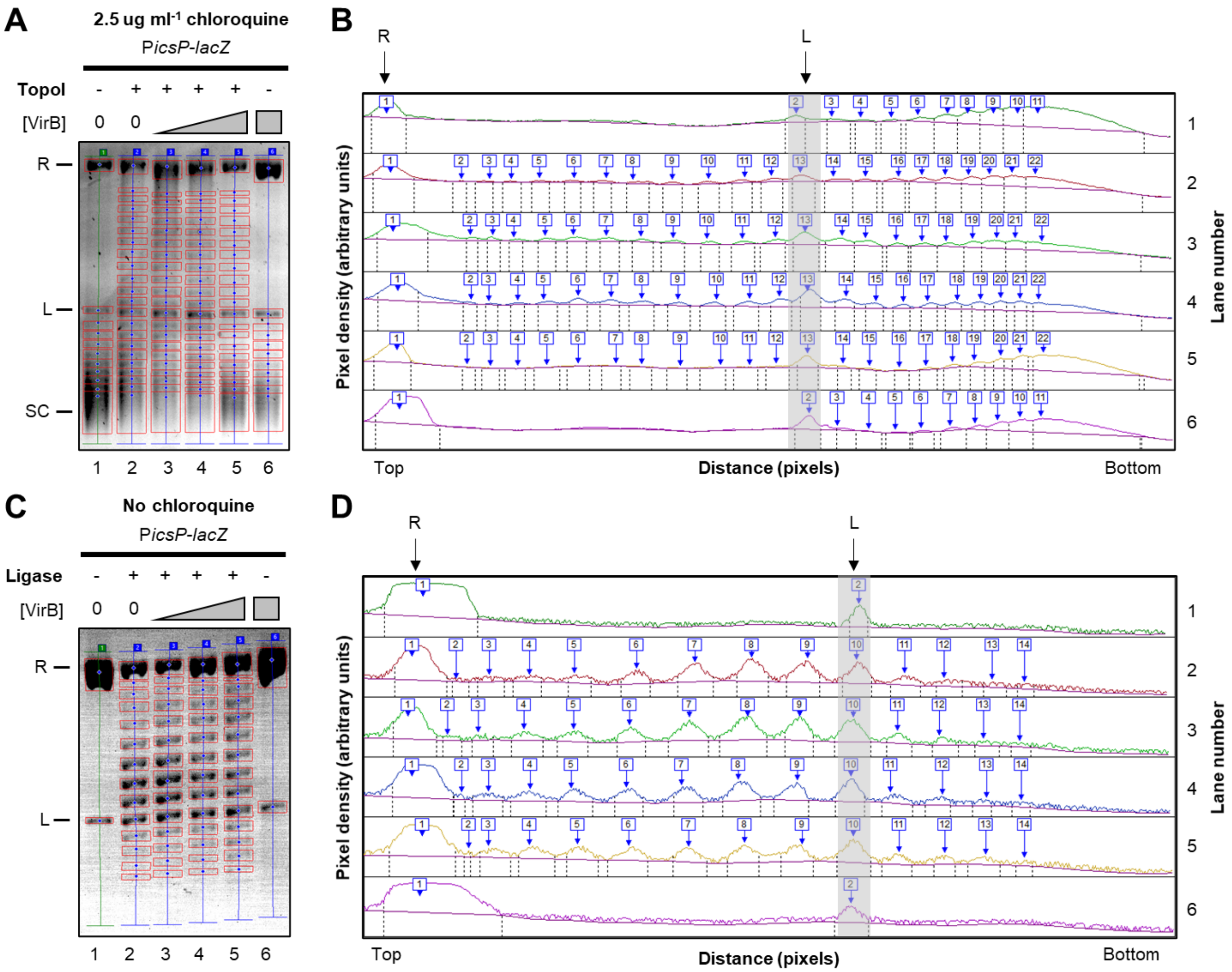


**Figure S8**: **Quantification of DNA incubated with VirB *in vitro* after 1D gel electrophoresis.** Quantification of images shown in Figure 6. A & C, show images of gel analysis. B & D, show stacked lane profiles (grey boxes, highlight linear bands denoted by L on both images, and R, represents the relaxed band). In both cases, VirB is seen in shift the topoisomer distribution up towards the top of the gel at low concentrations (compare lane 2 to 3) and with higher concentrations the topoisomer distribution drops down towards the bottom of the gel (compare lanes 3, 4 & 5). These changes in topoisomer distribution are consistent with VirB introducing positive supercoils into the DNA.
